# De Novo Design of Integrin α5β1 Modulating Proteins for Regenerative Medicine

**DOI:** 10.1101/2024.06.21.600123

**Authors:** Xinru Wang, Jordi Guillem-Marti, Saurav Kumar, David S. Lee, Daniel Cabrerizo-Aguado, Rachel Werther, Kevin Alexander Estrada Alamo, Yan Ting Zhao, Adam Nguyen, Irina Kopyeva, Buwei Huang, Jing Li, Yuxin Hao, Xinting Li, Aritza Brizuela-Velasco, Analisa Murray, Stacey Gerben, Anindya Roy, Cole A. DeForest, Timothy Springer, Hannele Ruohola-Baker, Jonathan A. Cooper, Melody G. Campbell, Jose Maria Manero, Maria-Pau Ginebra, David Baker

**Author notes:** Present address: Human Biology Division, Fred Hutchinson Cancer Center, Seattle, WA, USA. These authors contributed equally to this work (Xinru Wang, Jordi Guillem-Marti).

## Abstract

Integrin α5β1 is crucial for cell attachment and migration in development and tissue regeneration, and α5β1 binding proteins could have considerable utility in regenerative medicine and next-generation therapeutics. We use computational protein design to create de novo α5β1-specific modulating miniprotein binders, called NeoNectins, that bind to and stabilize the open state of α5β1. When immobilized onto titanium surfaces and throughout 3D hydrogels, the NeoNectins outperform native fibronectin and RGD peptide in enhancing cell attachment and spreading, and NeoNectin-grafted titanium implants outperformed fibronectin and RGD-grafted implants in animal models in promoting tissue integration and bone growth. NeoNectins should be broadly applicable for tissue engineering and biomedicine.

**One-Sentence Summary:** A de novo-designed fibronectin substitute, NeoNectin, is specific for integrin α5β1 and can be incorporated into biomaterials for regenerative medicine.

## Introduction

Integrin α5β1 is the principal receptor that directly binds to fibronectin (FN), a crucial extracellular matrix (ECM) component that is extensively expressed in various cells and tissues. The interactions between α5β1 and FN are vital for cell attachment and migration, making them integral to various stages of development and tissue regeneration, notably in wound healing, bone regeneration, and stem cell therapy^1–4^. However, the clinical use of full-length FN or its main interacting RGD (Arg-Gly-Asp) motif on biomaterials for regenerative purposes has been challenging. Full-length FN, typically derived from human plasma, poses challenges for large-scale production and is vulnerable to protease cleavage^5,6^. Conversely, while the RGD peptide can be easily manufactured and is widely used in biomaterial coatings, it does not elicit the desired cellular responses and does not consistently enhance bone formation in vivo^7–9^, perhaps because of the low affinity of RGD peptides to α5β1 compared to FN^10^ or because of the broad reactivity with the eight RGD/binding integrins^11,12^. The α5β1, α8β1, αvβ1, αvβ3, αvβ5, αvβ6, αvβ8, and αIIbβ3^13^ integrins all have the conserved RGD binding pocket with nearby glycan molecules making the design of integrin specific peptides challenging^11,12,14^(**Figures 1B-1F**); indeed to our knowledge there has been little success in developing high affinity integrin α5β1-specific binders.

**Figure 1:**
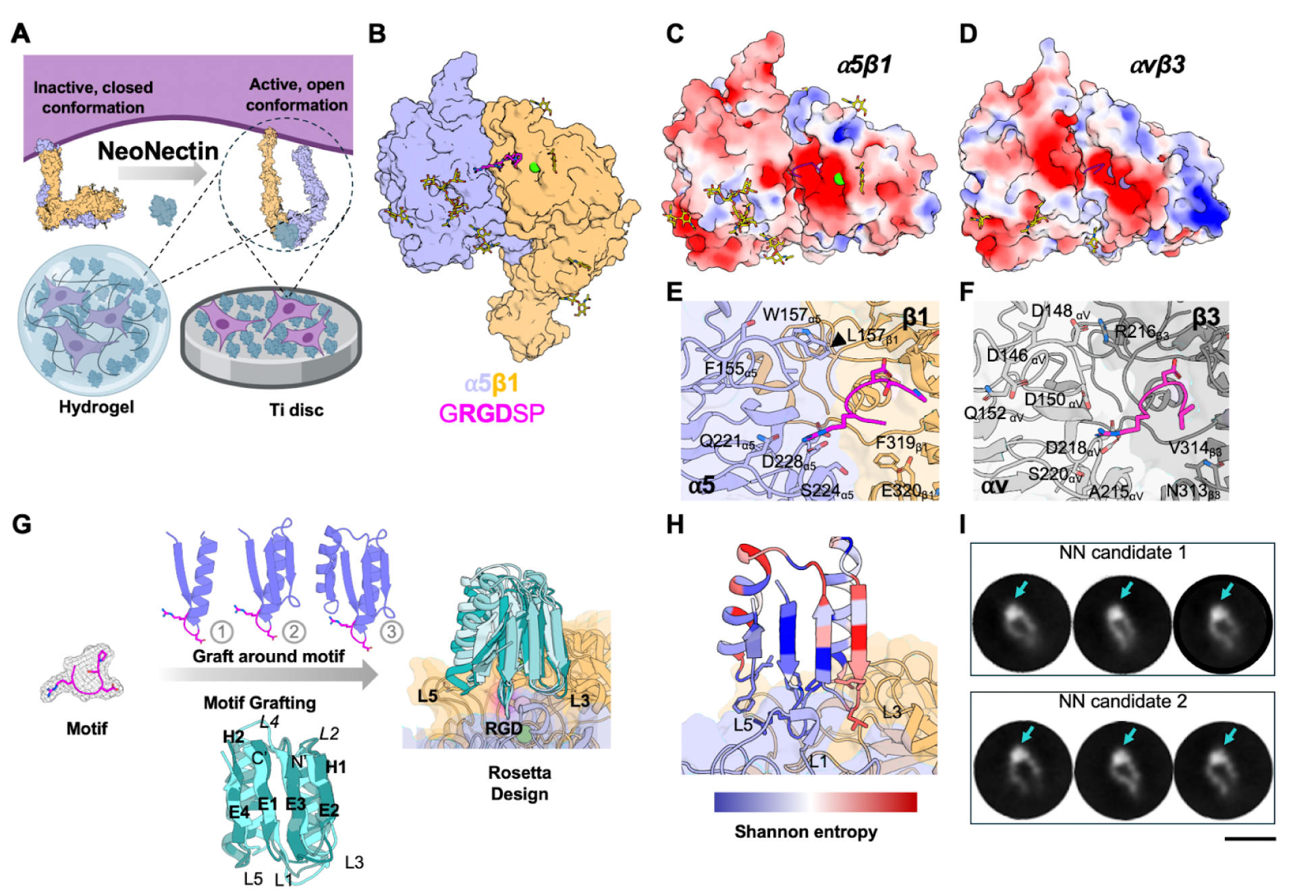
α5β1 binder design strategy. (A) Schematic of designed NeoNectin in biomaterial applications. (B) Previously solved crystal structure of RGD peptide (fibronectin^1492–1497^,magenta) bound α5β1(PDB: 4WK2) used for design. (C-D) Specificity design challenge highlighted by the similar electrostatic potential of integrins α5β1, αvβ3 (structures are from complexes with their cognate ligand peptides; PDB:4WK2 and 1L5G, respectively). Glycan molecules are shown as yellow sticks. (E-F) Zoomed-in views of RGD binding interfaces of α5β1 and αvβ3. (G) Design strategy for α5β1 specific NeoNectin. (H) Computational model of a designed α5β1 binder colored by site saturation mutagenesis results. The designed binder was colored by positional Shannon entropy, using a gradient from blue to red, with blue indicating positions of low entropy (conserved) and red those of high entropy (not conserved). (I) Representative negative stain electron microscopy 2D class averages showing NeoNectin candidates bind the open conformation of integrin α5β1 headpiece in buffer conditions (5 mM Ca^2+^) that do not favor FN binding. Each complex was formed by combining 1:2 molar ratio ⍺5β1:NeoNectin candidates. Arrows show additional density attributed to NeoNectin. Scale bar is 20 nm.

We reasoned that *de novo* protein design could enable the creation of small, stable, and easy to manufacture α5β1 binders that specifically activate α5β1 on biomaterials (**Figure 1A**). Integrin α5β1 undergoes a large conformational change from the inactive closed state to the active open state when bound to FN^15,16^. We set out to design α5β1 protein binders that can bind the interface between α5 and β1 subunits and stabilize the active conformation of α5β1. Due to their *in silico* origin and outstanding capacity to enhance cell adhesion, we call them NeoNectins (NN).

## Results

### Computational design

Following the design approach used to generate αvβ6 specific integrin binders^17^, we aimed to construct a protein scaffold capable of hosting multiple inserted loops to interact specifically with α5β1. Ferredoxin scaffolds, with their two helices and four beta strands, offer high stability and are ideal for loop presentation. Our goal was to utilize the three-loop side of the ferredoxin scaffolds to bind the groove formed between the α5 and β1 integrin subunits, targeting unique residues on α5 while stabilizing the open conformation of α5β1. We explored both the stepwise building up of ferredoxin starting from the RGD loop of FN extracted from an RGD/α5β1 complex structure (PDB:4WK2), or grafting the RGD loop onto computationally pre-built ferredoxin scaffold libraries (**Figure 1G**). In both cases, Rosetta flexible backbone protein design^18,19^ was then used to optimize the structure and the sequence of the design for shape complementarity and extensive interactions with α5β1. Designs were ranked based on Rosetta binding energy (ddG), solvent-accessible surface area, molecular contact surface, and a deep learning-based monomer folding metric^20^. A total of 7,820 designs from the first approach and 12,674 from the second approach were selected for experimental characterization.

### Experimental characterization

Synthetic oligonucleotides encoding the designs were cloned into a yeast surface-expression vector. Yeast cells displaying the designed proteins were incubated with biotinylated α5β1 ectodomain and several rounds of fluorescence-activated cell sorting (FACS) were used to enrich those that bound α5β1. The starting and enriched populations at each round were deep sequenced, and the frequency of each design in the starting population and after each sort was determined; this was used to estimate binding dissociation constants (K_D_) for each design^18^. For the 16 most enriched designs, we generated site saturation mutagenesis libraries (SSMs), in which every residue was substituted with each of the 20 amino acids, and sorted the SSMs in the presence of α5β1 at different concentrations and the affinity of each variant was calculated (**Figures 1H, S1A-S1F**). Substitutions at the RGD site in the SSMs were not tolerated for 8 out of 16 designs, as expected given the central role this motif plays in mediating integrin α5β1 and design interactions. Between 5-8 substitutions at designed interacting loop 3 and loop 5 that increased the apparent binding affinity were combined in small libraries which were sorted under stringent conditions (incubation with 0.2 nM biotinylated α5β1 followed by washing and overnight disassociation), yielding 20 optimized designs which were expressed in *E. coli* and purified (**Figures S1G and S1H**). Biolayer interferometry (BLI) showed that 5 designs bind to α5β1 with affinities ranging from subnanomolar to nanomolar K_D_ (**Figures 2A and S1I-S1L**).

**Figure 2:**
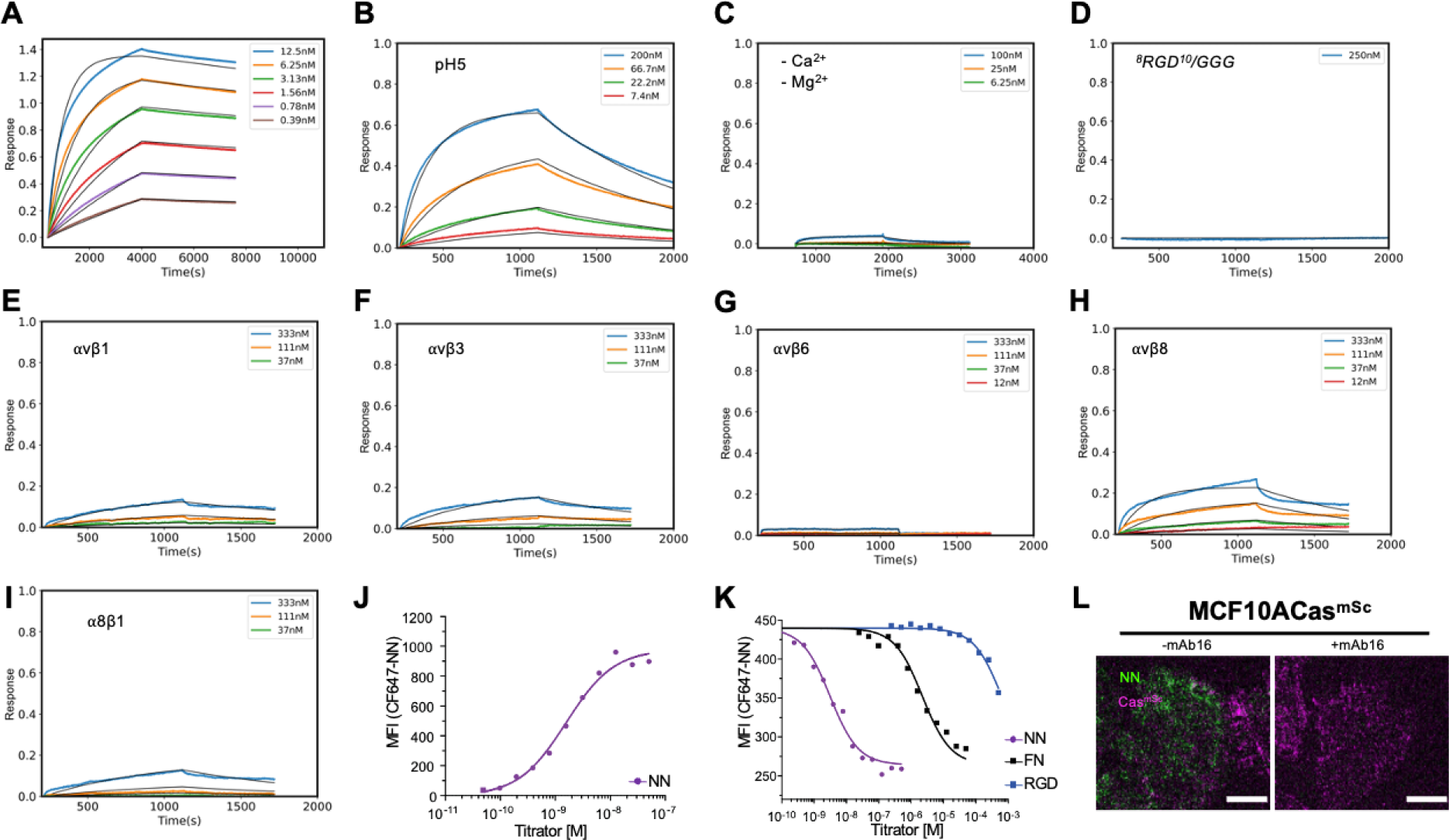
NeoNectin binds a5b1 with high affinity and specificity. (A-C) BLI binding affinity traces for NeoNectin against α5β1 in the resting buffer (20 mM Tris, pH = 7.4, 1 mM Ca^2+^, 1 mM Mg^2+^) or otherwise noted. Global kinetic fits, assuming a 1:1 binding model, were shown as black lines. (D) BLI binding affinity traces for NeoNectin ^8^RGD^10^/GGG mutant against α5β1. (E-I) NeoNectin does not bind integrins αvβ1, αvβ3, αvβ6, αvβ8, and α8β1. All affinity data was collected by Octet R8 and binding affinity was estimated by the Octet ForteBio software package. K_D_= 1.2 µM, 89 µM, can’t fitted, 1.0 µM and 0.32 µM, respectively. (J) Cell flow cytometry measurement for CF647 labeled NeoNectin against K562 cells, which only expresses α5β1 integrin. K_D_= 1.4 ± 0.3 nM. (K) Competing the binding of CF647 labeled NeoNectin by NeoNectin (NN), FN, RGD peptide on K562 cells gave KD values of 0.9 ± 0.2 nM, 612 ± 105 nM, 150000 ± 20000 nM respectively (see Methods for calculating KD values). (L) TIRF images of MCF10A cells treated with/without 200 nM α5β1-specific antibody mAb16 followed by addition of 20 nM GFP-tagged NeoNectin. Green: GFP-tagged NeoNectin. Each dot (in magenta) represents an integrin-positive structure marked by the presence of Cas protein. The representative images are of a single-cell image of MCF10A cells taken at 100X. The scale bar is 10 µm.

### NeoNectin binds to integrin α5β1 tightly and specifically

The design with the highest affinity binds to the integrin α5β1 ectodomain with a K_D_ of 0.4 nM at pH 7.4 and a K_D_ of 40 nM at pH 5 in the presence but not absence of metal cations measured by BLI (**Figures 2A-2C**). There was little binding towards other RGD binding integrins, measured by BLI or fluorescence polarization assays (**Figures 2E-2I, S2B-S2I**). Mutating the RGD motif to Gly-Gly-Gly (GGG) abolished binding (**Figure 2D**), supporting the importance of RGD motif in mediating integrin α5β1 binding. Circular dichroism melting studies showed that this design is stable up to 95 °C **(Figure S2A**). We refer to this highly selective and high affinity optimized design as NeoNectin throughout the remainder of the paper. To test if NeoNectin bound to α5β1 on cells, we made a CF647 (a cyanine-based far-red fluorescent dye) labeled NeoNectin and measured binding to wild-type K562 lymphoblast cells in which α5β1 is the only RGD-interacting integrin, obtaining an affinity of 1.4 ± 0.3 nM K_D_ (**Figure 2J**). Unlabeled NeoNectin competed CF647-labeled NeoNectin binding with a K_D_ within experimental error of 0.9 ± 0.2 nM. Competition with FN or RGD peptide showed much lower binding affinities, with IC50 values of 612 nM and 150000 nM, respectively (**Figure 2K**). Thus, NeoNectin binds to α5β1 680 times more tightly than FN and 16,700 times more tightly than RGD peptide.

We tested if NeoNectin binds cells expressing multiple RGD-interacting integrins in an α5β1-dependent manner using MCF10A Cas^mSc^ cells^21^ , a human mammary epithelial cell line expressing endogenously tagged Cas with mScarlet to mark the integrin positive adhesion^22,23^. Besides α5β1, MCF10A cell line also expresses other RGD binding integrins including αVβ1, αVβ3, αVβ6 ^24,25^. We first incubated MCF10A Cas^mSc^ cells^21^ in presence or absence of 200 nM α5β1-specific antibody mAb16 followed by incubation of 10 nM NeoNectin. The cells were then imaged by Total Internal Reflection Fluorescence (TIRF) microscopy. NeoNectin was only bound to MCF10A when cells were not pretreated with mAb16, suggesting it is specific for cellular α5β1 (**Figure 2L**). Because NeoNectin can bind to α5β1 tightly at low pH, we tested if it could be used to locate ligand-bound α5β1 in different cellular compartments. We observed colocalization of β1, NeoNectin, and endosome components RAB5, RAB11, and EEA19 (**Figures S3A-S3D)**, indicating a potential application in tracking α5β1 in cells with NeoNectin.

### NeoNectin-bound integrin α5β1 favors an extended open conformation

To investigate the effects of NeoNectin on α5β1 conformation and the molecular basis of the interactions, we used negative stain electron microscopy (nsEM) and cryogenic electron microscopy (cryo-EM). Many integrins, including α5β1 are known to undergo drastic conformational changes that are linked to activation state. In vitro, it has been established that high Ca^2+^ (5 mM) stabilizes the low affinity, closed headpiece conformation, while 1 mM Mn^2+^ stabilizes the high affinity conformation with an open headpiece and extended legs^26^. Using nsEM we found that in both cation conditions, NeoNectin binds and favors the extended-open conformation of integrin α5β1 (**Figure 1I, Figure S5E**).

Next, we determined the structure of α5β1 in complex with NeoNectin to a local resolution of 3.2 Å and an overall resolution of 3.3 Å (**Figures 3 and S4**). As expected, the dominant class (accounting for ∼95% of integrin particles) shows the α5β1 headpiece in an open conformation (**Figure 3, Figure S4A**) that is similar to the FN-bound integrin structure^16^. The interaction between NeoNectin and α5β1 in the experimentally determined cryo-EM model is consistent with the designed model in that it centers around three short loops: L1 (Gly7, Arg8, Gly9, Asp10, Phe11, Pro12), L3 (Asp33, His34, Lys35), and L5 (Gly58, Ile59, Trp60) (**Figures 3C-F**). The RGD motif encoded by L1 binds α5β1 via the same entities as the RGD motif in FN or RGD peptide^16,27^, specifically, α5 Gln221 and Asp227 bind Arg and β1-coordinated MIDAS cation coordinates Asp (**Figure 3D**). L3 further stabilizes the interaction through interactions with both subunits: His34 and Lys35 form a salt bridge triad with β1 Glu320 and His34 forms an additional interaction with α5 Ile225 (**Figure 3E**). FN also forms a salt bridge with β1 Glu320 that was suggested to be important for FN-induced headpiece opening^16^. However, interestingly when we mutated the positively charged His34 and Lys35 to Gly, we still observed the majority of particles in an open-headpiece conformation in activating and non-activating cation conditions. Our finding is in agreement with the ability of RGD peptides to open the integrin headpiece as shown by negative stain EM and by the ability of RGD peptides and FN to similarly stabilize the majority of the integrin conformational ensemble to the extended-open conformation^10^. NeoNectin L5 Trp60 interacts via shape complementarity with residues in both subunits, including a pi-pi stacking interaction with α5 Trp157. Trp157 has previously been shown to be the key for interacting with a Trp residue that introduces α5β1 specificity into RGD cyclic peptides; however, Trp157 is not required for binding to FN^12,16,28^. When mutated to Ala, the NeoNectin W60A variant still favors the open conformation of α5β1, thus suggesting it does not influence headpiece opening directly (**Figure S5G**). A list of experimentally determined interactions can be found in **Figure S5C**.

**Figure 3:**
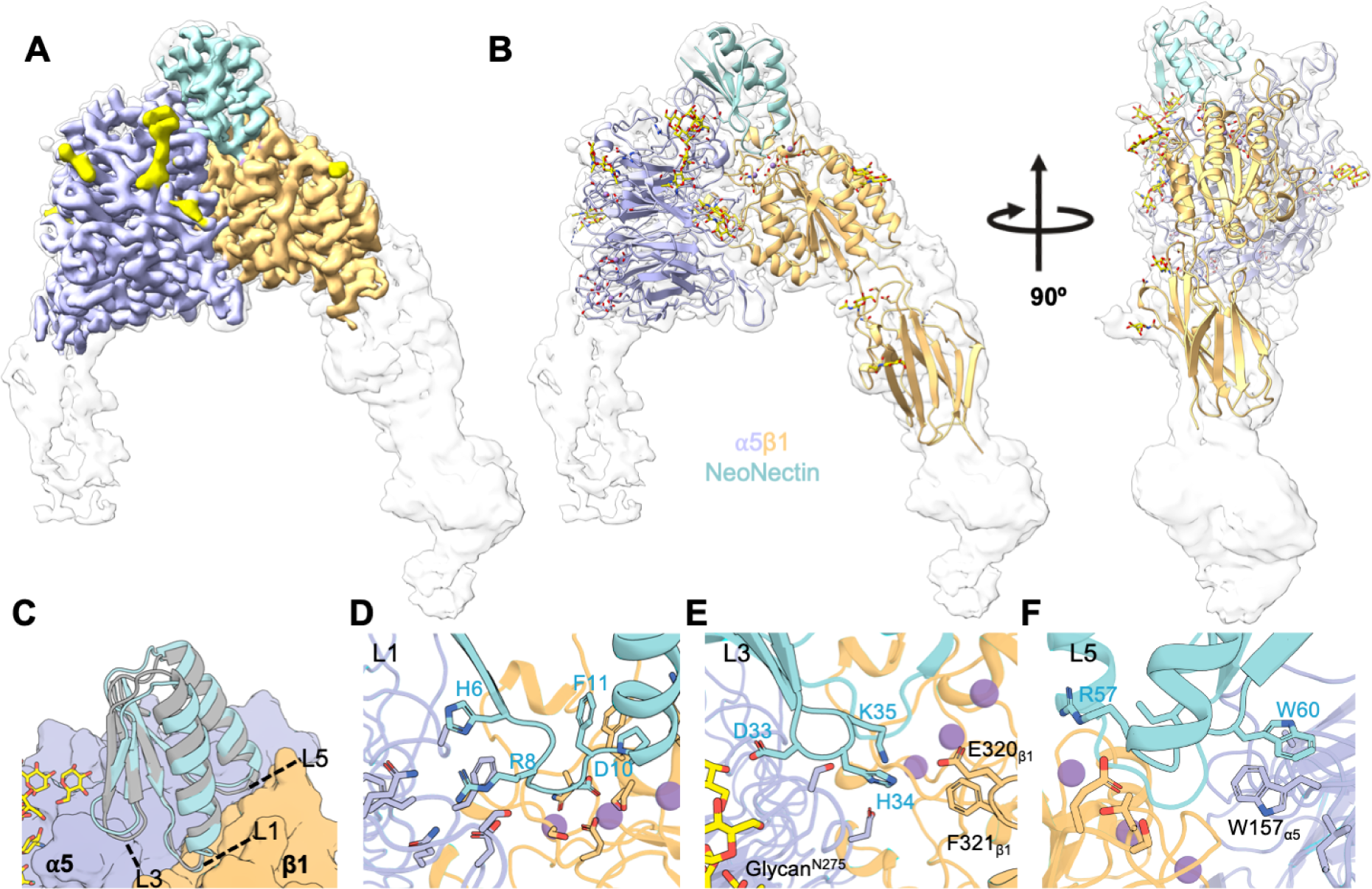
Structural characterization of integrin α5β1:NeoNectin complexes. (A) Cryo-EM map of α5β1 bound to NeoNectin. The sharpened, locally refined map is shown in color, superimposed with the unsharpened map in semi-transparent white. The color code is as follows: α5 (lavender), β1 (light orange), Neonectin (turquoise), coordinated cations (plum), glycans (yellow). (B) Two views of the ribbon model of α5β1 bound to NeoNectin displayed within the unsharpened density shown in (A). (C) An overlay of the NeoNectin-designed model (gray) and the experimentally determined model (turquoise). (D) Close-up of NeoNectin-L1(^6^HRGDFP^12^) and α5β1. R8 and D10 directly interact with α5β1 and other residues contribute to stabilizing the loop. (E) Close-up of NeoNectin-L3 (^33^DHK^35^) and α5β1 interface. (F) Close-up of NeoNectin-L5(^57^RGLW^60^) and α5β1 interface.

### Soluble NeoNectin inhibits α5β1-mediated cellular behaviors

Integrin α5β1 regulates cellular events such as cell attachment, cell migration, and tubulogenesis through interaction with FN found within the ECM. Both NeoNectin and FN bind to the RGD binding site of α5β1. Because of the high affinity and specificity of NeoNectin toward α5β1, we hypothesized that NeoNectin can inhibit α5β1-FN interaction dependent cellular responses as mentioned above. We first tested the effect of soluble NeoNectin on inhibiting spreading of MCF10A epithelial cells, which can migrate on ECM-containing FN and/or collagen^21^. We first plated MCF10A cells on either collagen- or FN-precoated glass surfaces and then treated the cells with NeoNectin at different concentrations for 30 min prior to washes. The remaining cells were imaged using fluorescent microscopy (**Figure 4A**). Soluble NeoNectin dramatically reduced cell attachment to FN-coated surfaces but not to collagen-coated surfaces, suggesting that NeoNectin does not interact with integrins specific for collagen-binding (**Figures 4B and 4C**). Similar effects were also observed in A549 adenocarcinoma human alveolar basal epithelial cells cultured on FN-grafted titanium discs or discs grafted with the cell attachment site (CAS) fragment from FN, but not laminin- (LAM, responsible for binding to integrins α1β1, α2β1, α3β1, α6β1, α7β1, α6β4 without involving RGD) and vitronectin-grafted (VTN; responsible for αvβ3 integrin binding) titanium discs **(Figures S6A and S6B**).

**Figure 4:**
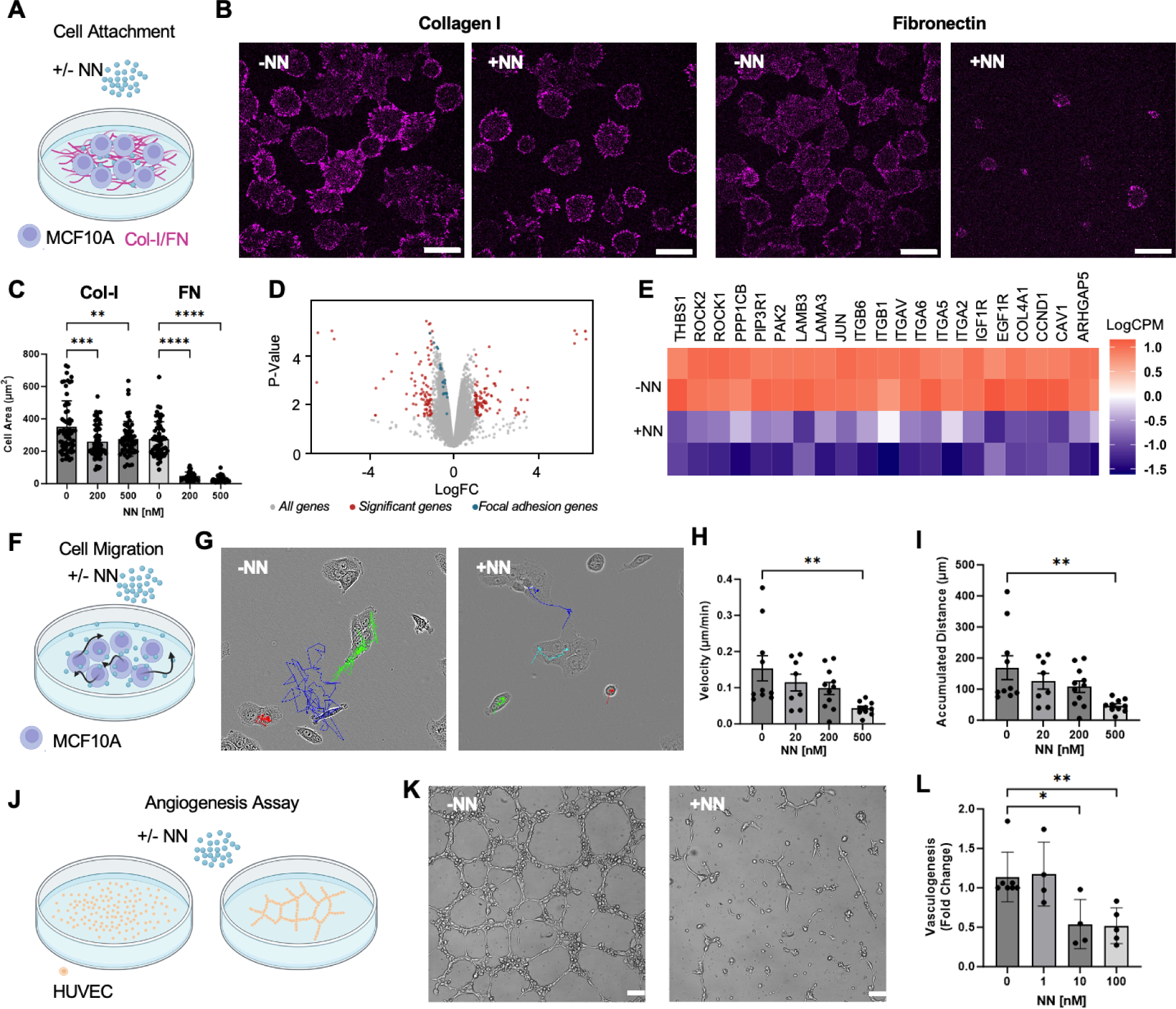
Soluble NeoNectin inhibits α5β1-mediated cell spreading and migration. (A) Schematic of the experimental design monitoring MCF10A cell attachment in presence of soluble NeoNectin on collagen I or FN coated surface. (B) Confocal imaging of MCF10A cells plated on collagen I or FN coated surface in presence of soluble NeoNectin for 30 minutes. The scale bar is 20 µm (C) Quantification of cell area in Figure 4B. (D) RNA-seq results from MCF10A cells plated on fibronectin surface in presence or absence of 500 nM NeoNectin for four hours, with genes related to the focal adhesion pathway highlighted in blue, genes significantly affected highlighted in red. (E) Heat map representation of genes involved in focal adhesion pathway. -NN: Cells spread on FN-coated surface. +NN: Cells spread on FN-coated surface in the presence of 500 nM NeoNectin. F, Schematic of the experimental design monitoring MCF10A cell migration with/without soluble NeoNectin. (G) Trajectories of individual cells tracked over an 18-hour imaging period in presence of 0 or 500nM NeoNectin. (H) Quantification of cell velocity in µm/min of individual cells from Figure G and Figure S6C. I, Quantification of accumulated traveled distance of individual cells from Figure G and Figure S6C. (K) Schematic of the tube formation assay. (L) Representative decrease in vascular stability by 10 nM soluble NeoNectin. Soluble NeoNectin were added to HUVEC cells at 0, 1, 10, and 100 nM. Vascular stability was analyzed after 12 hours. (M) The number of nodes, meshes, and tubes was quantified using an angiogenesis analyzer plug-in in ImageJ. The scale bar is 100 µm. Statistical significance was analyzed using One-way Anova Bonferroni’s multiple comparison test. All experiments have at least three biological replicates.

To investigate the effects of soluble NeoNectin on gene expression, we plated MCF10A cells on FN-coated plates and treated them with/without NeoNectin. After 4 h, we harvested the MCF10A cells and analyzed the transcript levels using RNA-Sequencing. Pathway enrichment analysis determined that NeoNectin treatment significantly down regulated focal adhesion-related genes (THBS1, ROCK2, ROCK1, PPP1CB, PIP3R1, PAK2, LAMB3, LAMA3, JUN, COL4A1, CCND1, CAV1, and ARHGAP5), integrin subunits (αv, α2, α5, α6, β1, and β6), and growth factor receptor tyrosine kinases (IGF1R and EGF1R) expression (p-value = 3.5x10^-7^) (**Figures 4D and 4E**).

Next we evaluated the effects of NeoNectin in inhibiting epithelial cell migration by culturing the cells on FN-coated surfaces and monitoring their positions every 10 min for 18 h after treatment with NeoNectin at different concentrations (**Figure 4F**). Cell migration is mediated by FN binding and recycling of integrins^29,30^, and hence, we expect the soluble NeoNectin to inhibit this process. Indeed, we observed a decrease in cell velocity and distance covered at all concentrations tested (**Figures 4H, 4I and S6C**).

We evaluated the capacity of NeoNectin to inhibit angiogenesis using human umbilical vein endothelial cells (HUVECs) in a tube formation assay (**Figure 4J**). Soluble NeoNectin was seeded with HUVECs at 0.1 to 100 nM or with PBS as control for 12 h before images were taken. Tube formation was significantly attenuated at 10 nM of NeoNectin (**Figures 4K and 4L**).

### Immobilized NeoNectin in hydrogel promotes cell attachment and spreading

Because NeoNectin binds integrin α5β1 more tightly than RGD and favors the extended-open conformation (**Figures 3A and 3B**), we hypothesized that NeoNectin could enhance cell attachment and cell spreading on biomaterials. We first investigated whether NeoNectin could enhance human mesenchymal stem cells (MSCs) adhesion and spreading in a 3D hydrogel (**Figure S7A**). Two NeoNectin variants with cysteines at solvent-exposed locations were (**Figure S7B;** NeoNectin^R25C^, and NeoNectin^E44C^) covalently tethered within poly(ethylene glycol) (PEG)-based material via the radical thiol-ene photopolymerization reaction^31,32,33^. MSCs encapsulated for 5 days with the NeoNectin^R25C^, and NeoNectin^E44C^ variants at a 0.5 or 1 mM concentration displayed significantly higher spread areas with robust stress fiber formation compared to the RGD condition, which displayed advanced protrusions, but very few stress fibers (**Figures 5A and 5B**); the cellular eccentricity was not statistically significant among the conditions (**Figure S7E**). At a concentration of 0.5 mM, MSCs encapsulated in the RGD condition remained rounded with very few protrusions, but those in the NeoNectin conditions were able to spread, resulting in significantly higher cell area and eccentricity, despite the lower availability of the adhesive moieties (**Figure S7C-S7G**). Thus, NeoNectin-modified PEG hydrogels effectively support the growth and function of encapsulated human cells.

**Figure 5:**
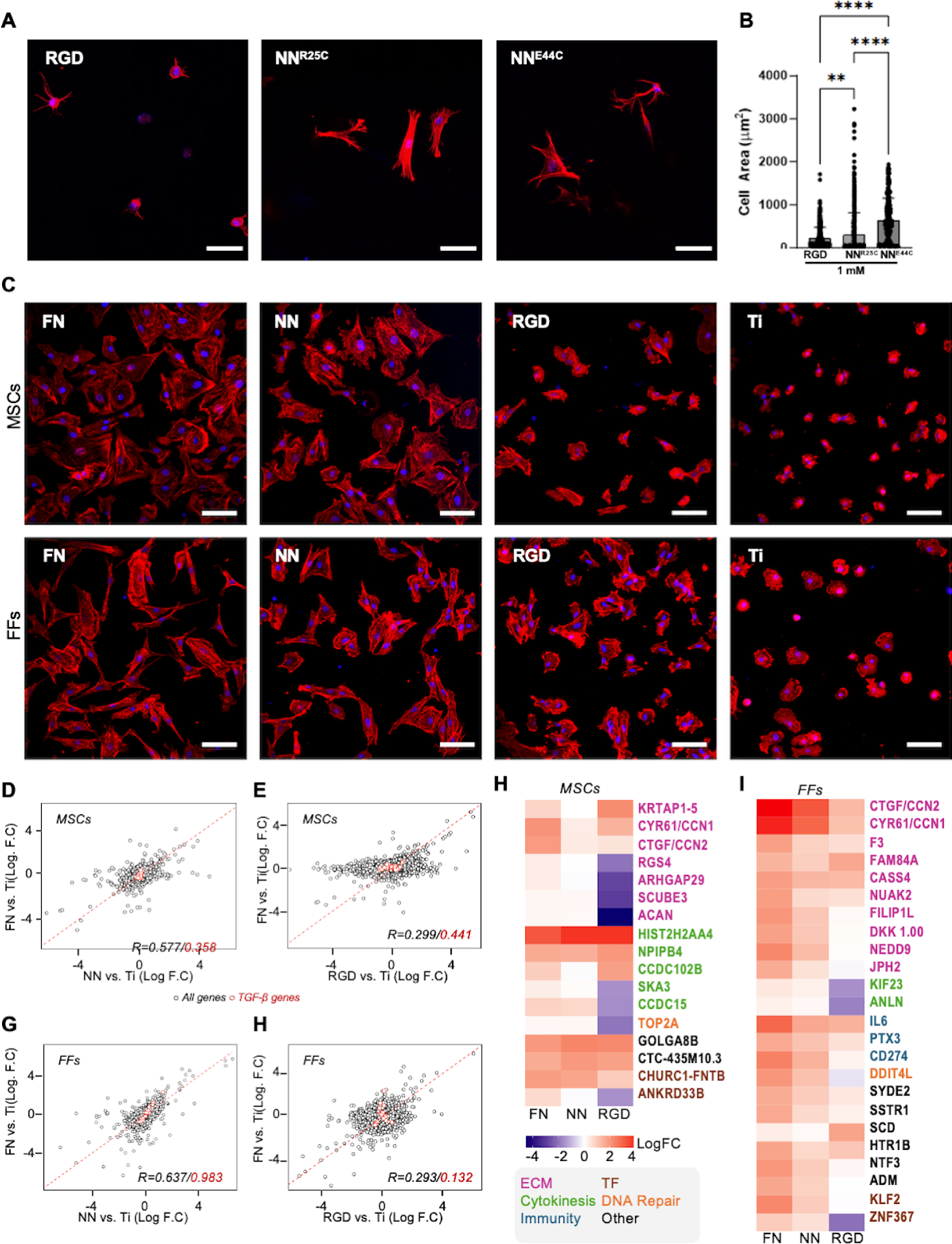
Immobilized NeoNectin stimulates FN-like cell spreading in 3D and 2D culture. (A) Representative immunofluorescence images of MSCs after 5 days of 3D culture within the different functionalized hydrogels. Adhesion modifications were included at 1 mM concentration. The scale bar denotes 50 µm. RGD: CRGDS. (B) Quantification of cell spread area in A. One Way ANOVA, Tukey’s Post-hoc Test. ** = p < 0.01, *** = p < 0.001, **** = p < 0.0001. (C) Representative immunofluorescence images of MSCs and FFs after 4 h of adhesion on the different functionalized surfaces. FN and NeoNectin were covalently linked through free amines. The RGD peptide (Cys-(Ahx)3-GRGDS) was covalently attached through the Cys. The scale bar denotes 100 µm. (D-G) Scatterplots of relative gene expression against bare Ti discs for MSCs or FFs between FN- and NeoNectin-grafted Ti discs. The genes of the TGF-β pathway for each cell type are highlighted in red circles. (H-I) Heat map representation of top differentially expressed genes compared to cells spread on bare titanium surface (LogFC > 1.5 or < -1.5 in MSCs). FN: Cells spread on fibronectin-grafted titanium surface. NN: Cells spread on NeoNectin-grafted titanium surface. RGD: Cells spread on RGD peptide-grafted titanium surface. ECM: Extracellular Matrix. TF: Transcription Factors.

### Immobilized NeoNectin on titanium discs promotes cell attachment and spreading

We then evaluated the behavior of bone tissue cells on NeoNectin-grafted titanium surfaces, one of the most commonly used materials in implantology. We analyzed the expression of different integrin subunits on different types of cells using qPCR, observing the highest expression of integrins α5 and β1 in MSCs and human foreskin fibroblasts (FFs), intermediate expression in human osteoblast (OBs) cells, and the lowest α5 and β1 expression in SaOS-2 and MG-63 osteosarcoma cell lines (**Figure S8A**).

MSCs cultured on NeoNectin-grafted titanium discs showed a completely spread morphology, similar to what we observed for MSCs cultured on FN-grafted surfaces (**Figures 5C and S8B**). Actin cytoskeletal filaments were well-developed and organized in both conditions (**Figure S8C**). Focal adhesions were also present, as indicated by vinculin staining, although to a lesser degree in NeoNectin-grafted surfaces compared to FN-grafted discs. The calculated area (size of each cell), the cellular circularity^34^ (the ratio of area to perimeter), and the number of cells (**Figure S8D**) attached to both surfaces presented no statistically significant differences. These data indicate that NeoNectin stimulates MSC attachment similarly to full-length FN, consistent with the expression of α5, αv, and β1 subunits (**Figure S8A**). In contrast, MSCs cultured on RGD-grafted titanium discs presented a round shape similar to bare titanium discs (**Figures 5C and S8B**), although with some protrusions and a diffuse actin cytoskeleton (**Figure S8C**). The area, the circularity and the number of cells in these two conditions were significantly lower than on NeoNectin- and FN-grafted surfaces (**Figure S8D**). Although MSCs express similar amounts of αv and α5 subunit, the high specificity of NeoNectin toward α5β1 suggests that integrin α5β1 is primarily responsible for MSC spreading.

FFs grown on FN- and NeoNectin-grafted titanium discs also showed similar morphology (**Figures 5C and S8B**), although fewer signs of focal adhesion were observed in the latter (**Figure S8C**). Similar to what we observed for MSCs, FFs grown on RGD-grafted and bare titanium discs presented a round morphology (**Figure 5C, S8B, S8C and S8D**). In contrast, other bone cells including OBs, and SaOS-2 and MG-63 cell lines grown on NeoNectin-grafted titanium discs presented a less spread morphology compared to FN-grafted surfaces (**Figures S8B and S8C**), in accordance with their lower levels of α5 and β1 integrins expression (**Figure S8A)**.

### Cells grown on NeoNectin- and FN-grafted titanium discs produce similar gene expression patterns

To test whether NeoNectin affects differential gene expression similarly to FN, we harvested MSCs, FFs, and OBs grown on FN-, NeoNectin-, RGD-grafted or bare titanium discs and performed bulk RNA-Sequencing. For both MSCs or FFs, we observed a strong correlation of gene expression when compared against bare titanium between FN and NeoNectin (R=0.577, R=0.637 respectively, **Figures 5D and 5E**) but not between FN and RGD (R=0.299, R=0.293 respectively,**Figures 5F and 5G**), suggesting that NeoNectin drives a transcriptional program more similar to FN than RGD. To determine if signaling was affected in a cell-type dependent manner, we further focused on genes involved in the TGF-β signaling pathway, an essential pathway involved in the activation of fibroblasts and the differentiation of MSCs/OBs^35,36^. Utilizing pathway enrichment analysis with Enrichr, we observe that FFs have upregulated TGF-β signaling in FN and NeoNectin conditions when compared to bare titanium (p-values: 5.7x10^-6^, 4.9x10^-4^ respectively), suggesting that both conditions have similar capabilities in signaling downstream in the TGF-β pathway as demonstrated by a strong correlation of magnitude of gene expression (R=0.984, **Figure 5F**). This result suggests that FN and NeoNectin activate FFs in a similar manner. In the case of MSCs and OBs, we find that TGF-β signaling is not significantly affected in all conditions (**Figures 5D, 5E, S9A and S9B**), suggesting that they maintain their differentiation potential.

To understand cell behavior differences, we compared the top differentially expressed genes relative to bare titanium. Genes with a log2 fold change greater or less than 1.5 were analyzed. Both MSCs and FFs grown on FN- and NeoNectin-grafted titanium discs showed increased expression of genes involved in ECM, cell attachment, proliferation, and survival (**Figures 5H and 5I**). Conversely, cells grown on RGD-grafted discs showed down-regulation of some of these genes. This is consistent with the less spread morphology of cells grown on RGD-grafted Ti discs or hydrogel (**Figures 5C, S8B, and S8C** ).

### NeoNectin-grafted implants enhance osseointegration

To evaluate the performance of the NeoNectin-grafted biomaterials *in vivo*, we quantified the osseointegration of implants in a rabbit cortical bone model. In brief, two implants were inserted per tibia of a rabbit (**Figure 6A**) so that each animal had all four conditions implanted (FN-, NeoNectin-, and RGD-grafted, and bare titanium). Samples were retrieved 3 and 6 weeks after implantation and the quality and degree of osseointegration for each sample were evaluated. In all conditions during the study, we observed no signs of infection or inflammation in the animals.

**Figure 6:**
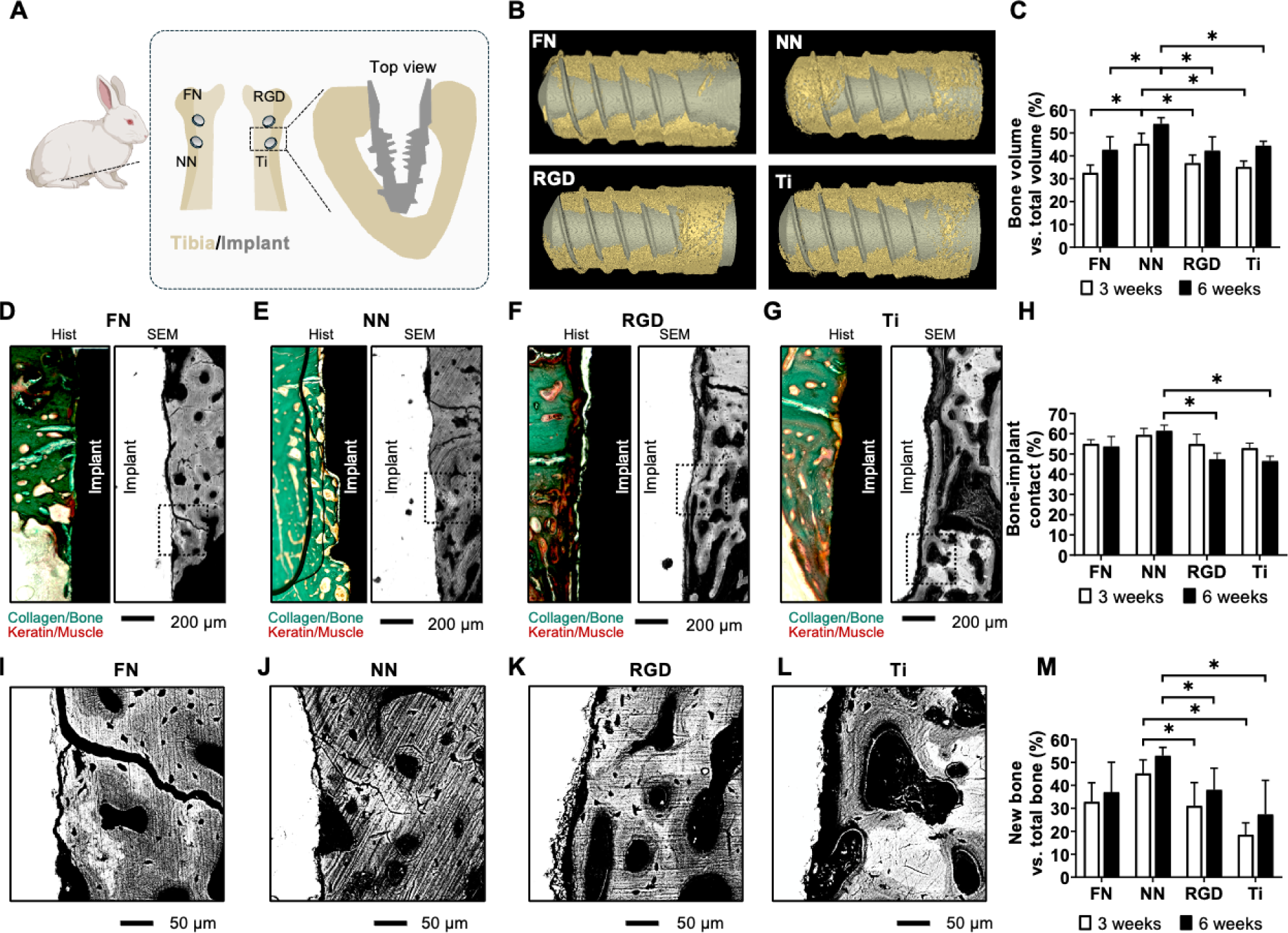
NeoNectin-grafted titanium implants outperform FN-grafted, RGD-grafted, and bare titanium implants in stimulating implant integration and bone growth. (A) Schematic of the in vivo experimental procedure with rabbits. Implants were randomly inserted into the tibia of rabbits, and samples were collected for histomorphometric analyses 3 and 6 weeks after the surgical intervention. N=7 for the 3 week and 6 week groups. (B) Representative micro-CT 3D reconstruction images showing bone (yellow) around the grafted or bare titanium implants (gray) 3 weeks post-surgery. (C) Calculated percentage of bone volume versus total volume (BV/TV) from micro-CT images. Non-parametric Mann Whitney’s test (* = p < 0.05). Data are presented as mean ± standard error of the mean. (D-G) Representative histological staining (left) and SEM (right) images of longitudinal sections 3 weeks post-implantation showing the implants conjugated with indicated molecules inserted into the tibia of rabbits. (H) Calculated bone-implant contact (BIC) percentage from SEM images. Non-parametric Mann Whitney’s test (* = p < 0.05). Data are presented as mean ± standard error of the mean. (I-L) Zoomed-in view of the boxed area in figures D-G. (N) Calculated percentage of new bone from the SEM images. Non-parametric Mann Whitney’s test (* = p < 0.05). Data are presented as mean ± standard error of the mean.

First, we evaluated the amount of bone around the implant. Extensive bone matrix was observed in micro-computerized tomography (micro-CT) 3D reconstructions (**Figures 6B and S10A**). The average ratio of bone volume to total volume (BV/TV) was the highest for NeoNectin in both the 3-week and 6-week groups (45.3% and 55.0%, respectively, **Figure 6C**). In contrast, FN, RGD, and titanium groups showed significantly less bone ratio. In all conditions, we observed an increase in the BV/TV ratio from 3 weeks to 6 weeks after implantation.

Next, we evaluated the quality of the bone around the implants. The bone around the NeoNectin-grafted implants appeared to be mostly compact and structured by histological staining and scanning electron microscopy (SEM) (**Figures 6D-6G and S10D-S10G)**. A more porous bone structure was noted at the interface of RGD-grafted and bare titanium implants (**Figures 6F-G and S10F-S10G)**, indicative of less mature bone. Furthermore, signs of fibrosis were mostly observed in RGD and bare titanium conditions by keratin red staining (**Figures 6F-G and S10F-S10G, left panels**). No signs of inflammation or infection were observed in the histological sections.

Then, we evaluated the bone integration by calculating the bone-implant contact (BIC) from SEM images. In all conditions, the implants were integrated into the cortical bone after 3 weeks with similar BIC ratios (**Figures 6D-G and S10D-G, right panels, 6H)**. However, all conditions except NeoNectin showed slightly decreased BIC values 6 weeks post-implantation. The most plausible explanation for this overall decrease in BIC values is that the growing bone was not interacting properly with the implants. This is evident in high magnification SEM images where new bone appears dark gray, while old bone is light gray (**Figure 6I-6L)**. After quantification (see example in **Figure S10B-S10C**), new bone around NeoNectin grafted implants showed the highest percentage relative to total bone at both 3 and 6 weeks of analysis (**Figure 6M),** suggesting that NeoNectin outperformed FN and RGD in promoting bone healing.

## Discussion

Most current biomaterials used for tissue regeneration, such as titanium or 3D hydrogels, do not promote sufficient cell attachment for successful tissue integration in transplanted tissues. Hence, there is a growing need for finding biomolecules that can be used for coating and embedding the biomaterials, making them more functional. Various levels of success have been achieved using RGD peptides or FN fragments, but their lack of specificity for various integrins and challenges in manufacturing constrain their application. Our designed NeoNectin shows considerable promise in overcoming these limitations. NeoNectin is a computationally designed 65 amino acid protein that is easy to produce at high yields and binds to integrin α5β1, favoring an extended-open conformation similar to FN bound α5β115,16. FN binds to both the α5-β1 interface and a synergy site on α5, while NeoNectin only binds to the interface between α5 and β1, suggesting that the synergy site binding is not essential for the conformational switching of integrin α5β1.

While several integrins are important in tissue regeneration, α5β1 is particularly crucial in the early stages for recruiting MSCs. However, RGD peptide-coated surfaces do not efficiently activate proper cell adhesion and spreading due to its lack of specificity and reduced affinity towards this integrin. Attempts to improve cell response have been made by cyclization of the RGD peptide, although α5β1 specificity has to our knowledge not been achieved. The high affinity and specificity of our designed NeoNectin for integrin α5β1 make it a promising candidate for immobilization into any biomaterial for tissue regeneration—cells grown on NeoNectin-grafted hydrogels or titanium implants completely spread and formed cytoskeletal fibers. NeoNectin remained active upon implantation of grafted titanium, demonstrating the potential for use in bone integration or other implantation applications.

We observed no inflammation or other undesired side effects with the recombinant protein-grafted material opening up the possibility of using computationally-designed proteins for regenerative medicine. Given the growing needs in this field, such as encapsulating therapeutic cells and/or recruiting/differentiating distinct cells during tissue healing, our approach of grafting materials with small, stable, target-specific designed proteins could be broadly applicable in regenerative medicine.

## STAR Methods

### Resource availability

#### Lead contact

Further information and requests for resources and reagents should be directed to and will be fulfilled by the lead contact, David Baker.

### Materials availability

This study did not generate new unique reagents.

### Data and code availability

● RNA-seq data have been deposited at GEO and will be publicly available on the date of publication. All the data reported in this paper will be shared by the lead contact upon request.
● All original code has been deposited at https://github.com/xinruwang7/NeoNectin and is publicly available as of the date of publication.
● Any additional information required to reanalyze the data reported in this paper is available from the lead contact upon request.
● The coordinates of the atomic models have been deposited in the Protein Data Bank under accession codes PDB: 9CKV.

### Experimental model and study participant details

#### Animals

A total of 20 mature New Zealand rabbit males were used in this study, each animal weighing 3.5-4 kg. All rabbits used were at 21-22 weeks of age. Ethical approval was obtained from the Ethics Committee of Consejería de Agricultura, Ganadería y Desarrollo Sostenible of the Junta de Extremadura (Spain) with ES 100370001499 authorization code (December 20, 2023). The rabbits underwent general anesthesia for the surgical procedures using an intramuscular mixture of dexmedetomidine (Dexdomitor; Ecuphar, Barcelona, Spain), ketamine (Ketamidor; Karizoo, Barcelona, Spain) and buprenorphine (Bupaq; Richter Farma, Wells, Austria). Anesthetic maintenance was performed by inhalation with isoflurane (Isoflo, Zoetis, Madrid, Spain) at a fixed concentration of 1-2%. In addition, Lidocaine at 20 mg/ml (Braun, Kronberg im Taunus, Germany) was administered via an infiltrative route in the operated area. A single incision was made on the internal region of each tibia in all animals. A full-thickness flap was opened, and randomized two implants were placed in the medial portion of each tibia near the epiphysis. Hence, four conditions were implanted in each animal. The implants were inserted according to the full-drilling protocol, with bicortical anchorage and separated by 6 mm in each tibia. A flat suture was made on the skin with simple stitches using 90% glycolide and 10% L-lactide 4/0 resorbable suture (Vicryl 4/0 Ethicon, Johnson & Johnson International, USA) to facilitate adequate primary wound closure.

### Method details

#### Computational design of integrin α5β1 binders

The ferredoxin scaffolds were generated in a piece-wise assembly manner. We first extracted the backbone information for all existing ferredoxin-like proteins reported by the SCOP database to generate guided blueprints for de novo ferredoxin. For the native ferredoxin-like fold with secondary structure “EHEEHE,” there existed four beta-sheets (E1, E2, E3, E4) and two helices (H1, H2) as basic elements, together with five loops (L1, L2, L3, L4, L5) connecting them in order. There existed optimal lengths and relative orientations between secondary structure elements. There also existed preferred torsion angles of the loops, which we further used abego to determine these loop torsion patterns. We identified 5 most occurring abego for each loop, 4 most occurring lengths for two helices (H1, H2), and 5 most occurring combinations of beta-sheets. Combining all possible variables, we finalized the top 100 blueprints encoding ferredoxin topology information for the next step. Based on the blueprints generated, we then applied the Rosetta blueprinter to create backbones constrained by the blueprints. To improve the success rate, we built each topology with 3 steps instead of constructing the whole protein in 1 step. We designed ferredoxin scaffold proteins either starting with the RGD peptide (Fibronectin^1492–1497^: 4WK2) or ferredoxin scaffolds library used for RGD-motif grafting in a later step. In the fast case step1, a helix and a beta strand were first built around the RGD peptide (E1+RGD+H1) with blueprint builder. After filtering with the Rosetta metrics, the top fragments were selected for step 2. In step 2, we further elongate a beta-hairpin at the C-terminus of the H1 and parallel to the E1 (E1+RGD+H1+L2+E2+L3+E3) and filtered the outputs with Rosetta monomer metrics. In step 3, the last L4+H2+L5+E4 fragments were built to make the whole ferredoxin. In the end, another round of filtering was imposed to remove designs with cavities and bad compactness. The trajectories that created the best-designed scaffolds were further resampled to increase the number of designs. The filtered backbones generated from the last step were sequence-optimized using Rosetta FastDesign and their structures were further evaluated via DeepAccNet^14^ or AlphaFold2^15^. Designs with plddt less than 0.85 were dropped. In the second case, we built the ferredoxin scaffolds libraries with altered steps.

#### Integrin α5β1 protein purification and biotinylation for yeast display

The integrin α5 (ITGA5, gene ID3678) β1( ITGB1, gene ID3688) ectodomain sequences were amplified by PCR reactions and inserted into the pD2529 vector. Integrin ectodomain were produced by co-transfecting α and β subunit cDNAs with C-terminal coiled-coils^37^ into Expi293F cells using FectoPro (Polyplus) according to the manufacturer’s instructions. The construct for the α5 subunit ectodomain in PD2529 CAG vector (ATUM) contains a N-terminal CD33 secretion peptide (MPLLLLLPLLWAGALA) and C-terminal HRV3C cleavage site (LEVLFQG), acid coil (AQCEKELQALEKENAQLEWELQALEKELAQ), Protein C tag (EDQVDPRLIDGK), and Strep twin tag (SAWSHPQFEKGGGSGGGGGSAWSHPQFEK). Construct for the β1 subunit ectodomain in PD2529 CAG vector contains an N-terminal CD33 secretion peptide and C-terminal HRV3C cleavage site, basic coil (AQCKKKLQALKKKNAQLKWKLQALKKKLAQ), HA tag (YPYDVPDYA), deca-histidine tag, P2A (ATNFSLLKQAGDVEENPGP), and mCherry. 24 hours of transfection, 3mM valproic acid and 4g/L of glucose were added. After 7 days of transfection, the integrin ectodomain was purified from the culture supernatant using His-Tag purification resin (Roche, cOmpelte^TM^, Cat No.5893682001), followed by size-exclusion chromatography (GE Healthcare, AKTA purifier, Superdex 200) as clasped ectodomain form. The purified protein was biotinylated with EZ-linkTM NHS-Biotin (catlog 20217, thermofisher) in 20 mM HEPES, pH7.4, 150 mM NaCl, 1 mM Ca^2+^, 1 mM Mg^2+^, with 10 µM α5β1 ectodomain and 100 µM protein and EZ-linkTM NHS-Biotin, at 37 degrees Celsius for 16 hours. Biotinylated α5β1 ectodomain was further purified by Superdex 200.

#### Integrin α5β1 protein expression and purification for BLI and structural determination

The integrin α5 (ITGA5, gene ID3678) β1( ITGB1, gene ID3688) ectodomain sequences were expressed in the pcDNA3.1-Hygro(-)-TET vector. The construct for the α5 subunit ectodomain contains a C-terminal HRV3C cleavage site (LEVLFQG), acid coil (AQCEKELQALEKENAQLEWELQALEKELAQ), and Strep-tag (WSHPQFEK). The construct for the β1 subunit ectodomain contains a C-terminal HRV3C cleavage site, base coil (AQCKKKLQALKKKNAQLKWKLQALKKKLAQ), and 6XHis tag. Integrin ectodomain was produced by co-transfecting α and β subunit plasmids containing C-terminal coiled-coils ^37^ into ExpiCHO cells (ThermoFisher) according to the manufacturer’s instructions. The integrin ectodomain was purified from the culture supernatant using a HisTrap Prepacked Column (Cytiva), followed by overnight protease cleavage and size-exclusion chromatography (GE Healthcare, AKTA purifier, Superdex 200).

#### Yeast surface display screening with FACS

The yeast surface display screening was performed using the protocol as previously described^14,16^. Briefly, DNAs encoding the minbinder sequences were transformed into EBY-100 yeast strain. The yeast cells were grown in CTUG medium and induced in SGCAA medium. After washing with integrin-FACS-buffer (20 mM Tris, 150 mM NaCl, 1 mM Ca^2+^, 1 mM Mg^2+^, and 1% BSA), the cells were incubated with 1uM biotinylated target proteins (integrin) together with streptavidin–phycoerythrin (SAPE, ThermoFisher, 1:100) and anti-c-Myc fluorescein isothiocyanate (FITC, Miltenyi Biotech, 6.8:100) for 60 min. After washing twice with integrin-FACS buffer, the yeast cells were then resuspended in the buffer and screened via FACS. Only cells with PE and FITC double-positive signals were sorted for next-round screening. After another round of enrichment, the cells were titrated with biotinylated target protein at different concentrations for 60 min, washed, and further stained with both streptavidin–phycoerythrin (SAPE, ThermoFisher) and anti-c-Myc fluorescein isothiocyanate (FITC, Miltenyi Biotech) at 1:100 ratio for 30 min. After washing twice with integrin-FACS buffer, the yeast cells at different concentrations were sorted individually via FACS and regrown for 2 days. Next, the cells from each subpool were lysated and their sequences were determined by next-generation sequencing.

#### Protein binder expression and purification

Synthetic genes encoding designed proteins were purchased from Genscript or Integrated DNA Technologies (IDT) in the pET29b expression vector or as eBlocks (IDT) and cloned into customized expression vectors^38^ using Golden Gate cloning. A His6x tag was included either at the N-terminus or the C-terminus as part of the expression vector. Proteins were expressed using autoinducing TBII media (Mpbio) supplemented with 50x5052, 20 mM MgSO_4_, and Studier trace metal mix in BL21 DE3 *E.coli* cells. Proteins were expressed under antibiotic selection at 25 degrees Celsius overnight after initial growth for 6-8 h at 37 degrees Celsius. Cells were harvested by centrifugation at 4000x g and resuspended in lysis buffer (20 mM Tris, 300 mM NaCl, 5 mM imidazole, pH 8.0) containing protease inhibitors (Thermo Scientific) and Bovine pancreas DNaseI (Sigma-Aldrich) before lysis by sonication. One millimolar of the reducing agent TCEP was included in the lysis buffer for designs with free cysteines. Proteins were purified by Immobilized Metal Affinity Chromatography. Cleared lysates were incubated with 0.1-2 mL nickel NTA beads (Qiagen) for 20-40 minutes before washing beads with 5-10 column volumes of lysis buffer, 5-10 column volumes of wash buffer (20 mM Tris, 300 mM NaCl, 30 mM imidazole, pH 8.0). Proteins were eluted with 1-4 mL of elution buffer (20 mM Tris, 300 mM NaCl, 300 mM imidazole, pH 8.0). All protein preparations were as a final step polished using size exclusion chromatography (SEC) on either Superdex 200 Increase 10/300GL or Superdex 75 Increase 10/300GL columns (Cytiva) using 20 mM Tris, 150 mM NaCl, pH 8.0. The reducing agent TCEP was included (0.5 mM final concentration) for designs with free cysteines. SDS-PAGE and LC/MS were used to verify peak fractions. Proteins were concentrated to concentrations between 0.5-10 mg/mL and stored at room temperature or flash frozen in liquid nitrogen for storage at -80. Thawing of flash-frozen aliquots was done at room temperature. All purification steps from IMAC were performed at ambient room temperature.

#### Enzymatic biotinylation of protein binders

Proteins with Avi-tags (GLNDIFEAQKIEWHE) were purified as described above and biotinylated in vitro using the BirA500 (Avidity, LLC) biotinylation kit. 840 µL of protein from an IMAC elution was biotinylated in a 1200 μL (final volume) reaction according to the manufacturer’s instructions. Biotinylation reactions were allowed to proceed at either 4 °C overnight or for 2-3 hours at room temperature on a rotating platform. Biotinylated proteins were purified using SEC on a Superdex 200 10/300 Increase GL (GE Healthcare) or S75 10/300 Increase GL (GE Healthcare) using SEC buffer (20 mM Tris pH 8.0, 100 mM NaCl).

#### Peptide Synthesis

Peptides were synthesized on a 0.1mmol scale via microwave-assisted solid-phase peptide synthesis (SPPS) LibertyBlue system (CEM) using preloaded Wang resin (CEM). The resin was subsequently treated with a cleavage cocktail consisting of TFA/TIPS/H2O/2,2-(ethylenedioxy)diethanethiol in 92.5/2.5/2.5/2.5 proportions for 3h, then precipitated in ice-cold ether and washed twice before drying under nitrogen. The resulting crude was resuspended in water and a minimal amount of acetonitrile and purified on a semi-preparative HPLC system (Agilent 1260 Infinity) with a linear gradient from solvent A to B of 2%/min (A: H2O with 0.1% TFA, B: acetonitrile (ACN) with 0.1% TFA). The peptide mass was confirmed via LC/MS-TOF (Agilent G6230B) and lyophilized to a white powder. For grafting titanium discs and implants, a long RGD peptide was synthesized in order to ensure accessibility to cells. This long RGD peptide consists of a 3-mercaptopropionic acid as anchoring moiety, 3 units of 6-aminohexanoic acid as spacer and the GRGDS sequence (MPA-(Ahx)_3_-GRGDS).

#### Circular Dichroism Spectroscopy

CD spectra were recorded in a 1 mm path length cuvette at a protein concentration between 0.3-0.5 mg/mL on a J-1500 instrument (Jasco). For temperature melts, data were recorded at 222 nm between 4 and 94 °C every 2 °C, and wavelength scans between 190 and 260 nm at 10 °C intervals starting from 4 °C. Experiments were performed in 20 mM Tris pH 8.0, 20 mM NaCl. The high tension (HT) voltage was monitored according to the manufacturer’s recommendation to ensure optimal signal-to-noise ratio for the wavelengths of interest.

#### Biolayer interferometry (BLI)

The BLI experiments were performed on an OctetRED96 BLI system (ForteBio) at room temperature in integrin resting buffer (20 mM Tris pH 7.4, 150 mM NaCl, 1 mM MgCl_2_, 1 mM CaCl_2_, 0.02 % Tween-20) or active buffer (20 mM Tris pH 7.4, 150 mM NaCl, 1 mM MnCl_2_,, 0.02 % Tween-20) or inactive buffer (20 mM Tris pH 7.4, 150 mM NaCl, 5 mM CaCl2, 0.02 % Tween-20) or low-pH buffer (20 mM Tris pH 5, 150 mM NaCl, 1 mM MgCl_2_, 1 mM CaCl_2_, 0.02 % Tween-20) Each BLI buffer was supplemented with 0.2 mg/mL bovine serum albumin (SigmaAldrich). Prior to measurements, streptavidin-coated biosensors were first equilibrated for at least 10 min in the assay buffer. Protein binders with N-terminal biotin were immobilized onto the biosensors by dipping them into a solution with 100 nM protein until the response reached between 10% and 50% of the maximum value followed by dipping sensors into fresh buffer to establish a baseline for 120 s. Titration experiments were performed at 25 degrees Celsius while rotating at 1000 rpm. Association of integrins was allowed by dipping biosensors in solutions containing designed protein diluted in octet buffer until equilibrium was approached followed by dissociation by dipping the biosensors into fresh buffer solution to monitor the dissociation kinetics. In the binding cross specificity assays each biotinylated binder was loaded onto streptavidin biosensors in equal amounts followed by 2 min of baseline equilibration. The association and dissociation with all the different binders were allowed for 900-1500 s for each step. Global kinetic or steady-state fits were performed on buffer-subtracted data using the manufacturer’s software (Data Analysis 12.1) assuming a 1:1 binding model.

#### Fluorescence Polarization

Binding affinity (or EC50) of the α5β1 binder to the soluble ectodomains of RGD-binding integrins by fluorescent polarization competitive binding assays. Affinities were measured by competing 10 nM FITC-cyclic-ACRGDGWCG (FITC labeled aminocaproic acid-disulfide-cyclized ACRGDGWCG peptide) binding to 200 nM αvβ1, 50 nM αvβ3, 50 nM αvβ5, 100 nM α5β1 or 1000 nM α8β1; 10 nM FITC-proTGFβ3 peptide (FITC labeled aminocaproic acid-GRGDLGRLKK peptide) binding to 10 nM αvβ6 or 250 nM αvβ8. In the assay, 10 µL of sample contains the 10 nM FITC-cRGD or FITC-proTGFβ3, integrin ectodomain, and binder with indicated concentrations was incubated at room temperature for 2 hours in the dark to ensure equilibrium before measurement. The buffer condition used for the reaction was 10 mM HEPES pH 7.5, 150 mM NaCl, 1 mM MgCl_2_, 1 mM CaCl_2_, and 0.5 mg/mL BSA. The competitive binding curves with binder or control as titrators on each individual integrin ectodomain were globally fitted with competitive binding equations^10^, with the maximum FP value in the absence of titrator and the minimum FP value when all the integrin in solution being bound by titrators as global fitting parameters, and K_D_ value for each titrator as individual fitting parameter. When K_D_ cannot be reliably fitted, the EC50 was calculated by fitting the curve with a three-parameter dose-response curve using Prism (GraphPad Software, version 9). The errors are the standard errors from the nonlinear least square fits.

#### Negative-stain EM sample preparation

The integrin-binder complexes were formed using a 1:2 integrin to NeoNectin molar ratio, incubated at room temperature for at least 10 min, and diluted to a final concentration of 90 μg/mL (α5β1) in 20 mM Tris-HCl pH 7.4, 150 mM NaCl, supplemented with either 1 mM MnCl_2_ or 5 mM CaCl_2_. 3 μL of the sample was applied to a glow-discharged 400 mesh copper glider grid that had been covered with a thin layer of continuous amorphous carbon. The grids were stained with a solution containing 2% (w/v) uranyl formate as previously described^39^.

#### Negative-stain EM data acquisition and processing

Data were acquired using a Thermo Fisher Scientific Talos L120C transmission electron microscope operating at 200 kV and recorded on a 4k × 4k Thermo Fisher Scientific Ceta camera at a nominal magnification of 92,000× with a pixel size of 0.158 nm. Leginon was used to collect >100 α5β1 micrographs at a nominal range of 1.8–2.2 μm under focus and a dose of approximately 50 e^−^/Å^2^. Experimental data were processed using Gautomatch (https://github.com/JackZhang-Lab), RELION, and cryoSPARC^40–42^.

#### Cryo-EM sample preparation

The integrin-binder complexes were incubated at room temperature for 30 min using a 1:2 integrin to binder molar ratio. From there, complexes were diluted to a final concentration of 90 µg/mL (α5β1) in 20 mM Tris pH 7.4, 150 mM NaCl, 1 mM MnCl_2_. For cryo-EM grid preparation, UltrAufoil grids (300 mesh, 1.2/1.3) were glow-discharged for 30 s at 15 mA, then 3 μL of each complex were applied to each grid. Complexes were frozen with a Thermo Fisher Scientific Vitrobot Mark IV in 100% humidity at 4 °C and vitrified in liquid ethane cooled by liquid nitrogen.

#### Cryo-EM data acquisition, processing, and model building

Datasets were acquired on a Thermo Fisher Scientific Glacios cryo-transmission electron microscope operating at 200kV and recorded with a Gatan K3 Direct Detection Camera. A total of three datasets were collected from three separate grids. For data collection, the stage was tilted to 30° and all images were recorded using SerialEM software^43^. One hundred frame movies were recorded in super-resolution mode with a super-resolution pixel size of 0.561 Å/px, a nominal magnification of 36kx, a nominal defocus range of 1.2 to 2.0 μm under focus and an approximate dose of 50 e^−^/Å^2^. 861 micrographs were used for subsequent data analysis (334, 183, 204 micrographs from the respective data collections). Dose fractionated super-resolution image stacks were motion corrected and binned 2 × 2 using Fourier cropping with MotionCor2 within the RELION wrapper^44^. Motion corrected stacks were processed using Patch CTF in cryoSPARC. Initially, 728,379 particles were picked using the unbiased blob picker in cryoSPARC and subjected to iterative 2D and 3D alignment and classification yielding a final map at 3.28Å resolution (72,604 particles). Additional local refinement of the binding interface gave a final map at an improved resolution of 3.19Å. The resulting orientational distribution and gold-standard Fourier shell correlation plots are shown in **Figure S4**. Model building: An open α5β1 headpiece structure (PDB: 7NWL) and the designed binder model were used as initial models. First, these models were fit into their respective cryo-EM density using UCSF ChimeraX and glycans were manually added. Refinements were performed using COOT and ISOLDE^45,46^. All maps (sharpened and unsharpened) used for modeling have been deposited. Cryo-EM and model building statistics can be found in **Table S1**.

#### Mammalian cell culture

Human umbilical vein endothelial cells (HUVECs) were purchased from Lonza (Catalog #: C2519AS) and cultured in EGM2 media as described previously (Zhao et al 2021). HUVECs were expanded and serially passaged to reach passage 4 before cryopreservation.

MCF10A cells were cultured in media described previously^50,51^; briefly, the media consisted of DMEM/F12 (Gibco, 11330032), 5% horse serum (Gibco, 16050130), 20 ng/mL EGF (Sigma-Aldrich, SRP3027), 0.5 mg/mL hydrocortisone (Sigma-Aldrich, H4001), 100 ng/mL cholera toxin (Millipore, C8052), 10 µg/mL insulin (Sigma-Aldrich,11070-73-8) and 1% penicillin–streptomycin (Gibco, 15140122). MCF10A cells were starved in the same media without EGF and contained 2% horse serum (assay media) for 16 hours before signaling experiments.

Human bone marrow-derived mesenchymal stem cells (MSCs; ATCC) and human osteoblasts (OBs; Sigma-Aldrich) were cultured in Advanced DMEM medium supplemented with 10% FBS, 20 mM HEPES, penicillin/streptomycin (50 U/mL and 50 µg/mL, respectively) and 2 mM L-glutamine (all components from ThermoFisher Scientific). Cells from passage 5 were used in all experiments.

Human foreskin fibroblasts (FFs; Millipore), MG-63 cells (ATCC), and A549 cells (Elabscience) were cultured in DMEM medium supplemented with 10% FBS, 20 mM HEPES, 50 U mL^-1^ penicillin, 50 µg mL^-1^ streptomycin and 2 mM L-glutamine, all from ThermoFisher Scientific. FFs from passage 10 were used in all experiments.

SaOS-2 cells (ATCC) were cultured in McCoy’s 5A medium supplemented with 10% FBS, 20 mM HEPES, 50 U mL^-1^ penicillin, 50 µg mL^-1^ streptomycin and 2 mM L-glutamine, all from ThermoFisher Scientific.

#### Cell binding assays using flow cytometry

CF647 Succinimidyl Ester (Biotium 92135) was used to directly label NeoNectin following the manufacturer’s protocol. To determine the affinity of CF647- NeoNectin to α5β1 on the K562 cell surface (**Figure 2J**), 100 µL of cells (10^6^/mL) were mixed with indicated concentrations of CF647- NeoNectin L15 medium containing 1% BSA for 2 hrs at room temperature and subjected to flow cytometry without washing. Background fluorescence was measured with 10 mM EDTA in the binding buffer. The background-subtracted mean fluorescence intensity (MFI) at each concentration of CF647- NeoNectin was fitted to a three-parameter dose-response curve for K_D_, background MFI, and maximum MFI.

Affinities of unlabeled NeoNectin, FN fragment (Fn3_9-10_) and RGD peptide (GRRGDGATGH) for intact α5β1 on K562 cells were measured by competing CF647- NeoNectin binding (**Figure 2K**). Cells (10^6^/mL in 100 µL) were mixed with 5nM CF647- NeoNectin and the indicated concentrations of competitor in L15 medium with 1% BSA. After 2 hrs in the dark at room temperature to ensure equilibrium, cells were subjected to flow cytometry without washing. MFI of CF647- NeoNectin at each concentration of different competitors were globally fitted to three parameter dose-response curves, with maximum MFI in absence of competitor and minimum background MFI as shared fitting parameters, and EC50 value for each competitor as individual fitting parameter. With the fitted EC50 value, K_D_ of each competitor was calculated as K_D_ = EC50 / (1+ C_L_/K_D, L_), where C_L_ is the concentration of CF647- NeoNectin used (5 nM), and K_D, L_ is the binding affinity of CF647- NeoNectin to α5β1 determined in **Figure 2J**.

#### Covalently grafting titanium surfaces with different biologics

NeoNectin and FN were covalently immobilized through their primary amine groups as previously described for other proteins^47,48^. Briefly, Ti discs of 10 mm diameter and 2 mm thickness were polished to remove the effect of roughness on cell attachment using silicon carbide papers and colloidal silica. Afterward, the discs were ultrasonically rinsed with cyclohexane, isopropanol, deionized water, ethanol, and acetone. Then, Ti discs were activated by oxygen plasma at 12 MHz in a Femto low-pressure plasma system (Diener Electronic) and immersed in a 0.08 M solution of (3-aminopropyl)triethoxysilane (APTES, Sigma-Aldrich) at 70 °C for 1h, rinsed with different solvents, and APTES was cross-linked with 7.5 mM solution of N-succinimidyl-3-maleimidopropionic acid. NeoNectin was then added at a 100 µg/mL (10 µM) concentration in PBS, being covalently linked through the amines of exposed lysines. FN was added at a 50 µg/mL concentration and covalently attached through its free amines. The RGD peptide (MPA-(Ahx)_3_-GRGDS) was chemically synthesized as described above and added at a 100 µM concentration in PBS at pH 6.5, and covalently attached through the free thiol.

#### Cell adhesion and spreading assay on titanium surface

MSCs, FFs, OBs, SaOS-2, and MG-63 were cultured in serum-free conditions at a concentration of 25,000 cells/disc and allowed to adhere for 4 h on the grafted titanium discs. Then, cells were fixed with 4% paraformaldehyde for 20 min and rinsed with 20 mM glycine in PBS (washing buffer). Cells were permeabilized with 0.05% Triton X-100 in PBS for 15 min, rinsed thrice with washing buffer, and blocked with 1% BSA for at least 30 min. Cells were then incubated with mouse anti-vinculin (1:100; Sigma-Aldrich) for 1h, rinsed with washing buffer, and incubated with Alexa Fluor 488 goat anti-mouse (1:1000; ThermoFisher Scientific) and Alexa Fluor 546 phalloidin (1:300; ThermoFisher Scientific) for 1 h in the dark. Samples were mounted in a mounting medium containing DAPI for counterstaining the nuclei and visualized in an LSM 800 confocal microscope (Carl Zeiss). The area of cells and the circularity were calculated using the ImageJ software in images from at least five areas randomly selected.

#### Tube formation assay

Tube formation assay was performed using previously described protocol^49^. Briefly, passage 4 HUVECs were thawed onto a 10 cm 0.1 % gelatin pre-coated plate and cultured until 80-90% confluent. Before cell seeding, 150 μL of 100% matrigel is added to a pre-chilled 24-well plate to allow even spreading of the matrigel. Allow the matrigel plate to solidify at room temperature for 25 min. Seed HUVECs at 150,000 cells per 350 μL in each well with PBS or NeoNectin at 0.1 to 1000 nM, considering 500 μL as the total volume in each well. Cells were then imaged after 12 hours. 20 images were taken in each well at random locations and images were analyzed using the Angiogenesis analyzer plugin in ImageJ. An average of the number of nodes, meshes, and segments of the 20 images, and these three parameters were also averaged to calculate the vascular stability for each well.

#### Cell spreading assays and microscopy

The glass bottom dishes (FluoroDish, FD35-100, World Precision Instruments) were precoated with 50 μg/mL collagen-I (Advance Biomatrix, #5056) or 5 μg/mL FN (Sigma, #F1141) for at least 3 h at 37 °C and washed with PBS. Integrin binder was added as indicated concentration on the precoated glass-bottom dish for 30 min and cells were plated on these surfaces for an additional 30 min before fixing them with 4% paraformaldehyde (PFA). The PFA fixed cells were permeabilized with the help of 0.1% Triton-X-100. Cells were blocked with 2% BSA, and 5% normal goat serum in PBS for 30 min followed by three washes with 1XPBS and incubated for 1 h with Alexa Fluor 647 Phalloidin (1:100 dilution) (Thermo, #A22287) for F-actin staining. The images were taken after washing thrice with 1XPBS on an automated TIRF microscope (Nikon Ti, 100x/1.49 CFI Apo TIRF oil immersion objective) equipped with Perfect Focus, motorized x-y stage, fast piezo z stage, and Andor iXon X3 EMCCD camera with 512 x 512-pixel chip (16-micron pixels). These images were processed and analyzed using ImageJ.

#### Single-cell migration inhibition assays

MCF10A cells (5X 10^3^) were plated onto 12 wells plate in assay media. These cells were treated with the increasing concentration (0, 20, 200, 500 nM) of integrin mini binder (mb) after 12 h of attachment and imaged once every 10 min for 18 h on an IncuCyte S3. Images were processed using ImageJ software and analyzed using a manual tracking plugin.

#### RNA-Sequencing

MCF10A cells were prepared as described in cell adhesion inhibition assays. MSCs and FFs were prepared as described in cell attachment and spreading experiments. 5x10^4^ –1x10^5^ cells were harvested. RNA was prepared by directly lysing cells in plates with 350 µL RLT Plus with B-Me and processed with the RNAEasy Plus mini kit (Qiagen cat. no 74134) to obtain gDNA-eliminated total RNA. RNA was further processed into bulk RNA-seq libraries (1 ug input per library) in duplicate with an Illumina Stranded mRNA prep kit (Illumina cat. no 20040532) according to the manufacturer’s instructions. Final libraries were quantified and characterized with an Agilent High Sensitivity D1000 ScreenTape (Agilent cat. no 50675584). Libraries were sequenced on an Illumina P2 100 cycle kit with the following parameters: 10:59:59:10 (index1:read1:read2:index2). Data was demultiplexed with bcl2fastq and preprocessed with BioJupies. The resulting count matrices were log CPM normalized per sample and z-scored across conditions to compare expression levels. Limma was used to compare DEGs. Enrichr was used for GSEAs.

#### Hydrogel

The dicysteine crosslinking peptide Ac-GCRDLPESGGPQGIWGQDRCG-NH_2_ was purchased from Genscript (Piscataway, NJ). The FN-derived adhesion sequence CRGDS was synthesized on rink amide ProTide resin (CEM Corporation; Charlotte, NC) following induction-heating assisted Fmoc solid-phase techniques with HCTU activation (Gyros Protein Technologies PurePep Chorus; Tucson, AZ) at a 0.2 mmol scale. The resin was treated with a trifluoroacetic acid (TFA)/ethane dithiol (EDT)/water/triisopropylsilane (94:2.5:2.5:1) mixture for 3 hours, then precipitated and washed in ice-cold diethyl ether (2 x 150 mL). The crude peptide was purified via semi-preparative reversed-phase high performance liquid chromatography with a linear gradient of 5-100% acetonitrile and 0.1% TFA for 45 minutes and then lyophilized to yield a white powder of the final peptide CRGDS. Peptide mass was verified via ESI-LCMS. Both peptides were resuspended in 10% acetic acid and lyophilized to yield aliquots of the desired mass.

For MSC encapsulation, all gel precursors were combined at 3 mM 4-arm Poly(ethylene glycol) norbornene terminated (PEG-NB; JenKem): 12 mM dicysteine peptide: 1 mM lithium phenyl-2,4,6-trimethylbenzoylphosphinate (LAP; Allevi3D). RGD peptide (CRGDS) or the NeoNectin variants were included in the final formulation at a concentration of either 0.5 or 1 mM. MSCs were resuspended in the gel mixture at a concentration of 1x10^6^ cells/mL and 5 µL gels were pipetted on the bottom of a 96 well-plate, at which point they were exposed to collimated near-UV light (λ = 365 nm; 10 mW cm^-2^; Omnicure 1500) for 2 minutes to allow for thiol-ene polymerization. The gels were then covered in media and cultured for 5 days. On day 5, gels were fixed by treatment with 4% paraformaldehyde for 1 hr at room temperature, and then washed 3 x 10 minutes with PBS and permeabilized for 30 minutes with 0.5% Triton X-100 in PBS. Subsequently, actin was labeled with 1:400 Phalloidin AF-532, and nuclei—with 1:1000 Hoechst 33342 in PBS. Gels were rinsed in PBS and imaged on a Leica Stellaris confocal microscope. Cell area and eccentricity were analyzed with Cell Profiler 4.0^50^.

#### Colocalization imaging

MCF10A Cells were plated on 35 mm glass-bottom dishes (FluoroDish, FD35-100, World Precision Instruments) for 24 h. These cells were treated with C6-GFP-NN for 30 min and incubated at 37°C containing 5% CO_2_. Cells were fixed with 4% paraformaldehyde (PFA) and permeabilized with 0.1% Triton-X-100. The non-specific antigens were blocked with blocking reagents (2% BSA+3% normal goat serum), which was followed by incubation with primary antibodies for 2 h. The antibodies used in the study are rat anti-integrin-β1 (9EG7, 553715) mouse anti-Rab-5 (BD Transduction Laboratories, 610724), mouse anti-Rab-11 (BD Transduction Laboratories, 610656), mouse anti-EEA-1 (BD Transduction Laboratories, 610456). The cells were washed with 1X PBS thrice before adding appropriate secondary antibodies. These cells were mounted in ProLong Gold (Invitrogen) for confocal microscopy using a Dragonfly 200 High-speed Spinning disk confocal imaging platform (Andor Technology Ltd) on a Leica DMi8 microscope stand equipped with a ×100/1.4 oil immersion objective, iXon EMCCD and sCMOS Zyla cameras and Fusion Version 2.3.0.36 (Oxford Instruments) software together with Imaris simultaneous deconvolution. These images were used to evaluate the percentage of colocalization using Mander’s colocalization coefficient between Rab5 and integrin binder with ImageJ plugin Coloc 2.

#### Animal implantation

Titanium implants of 3 mm diameter and 8 mm length with a conventional sandblasted, large grit, acid-etched (SLA) surface (Klockner Vega implants; Soadco, Andorra) were functionalized as described above for titanium discs. Four conditions were prepared: (i) FN, (ii) NeoNectin, (iii) MPA-(Ahx)3-GRGDS (RGD peptide) (iv) bare Ti. After 3 and 6 weeks, animals were euthanized (8 animals per time) using the same anesthesia protocol mentioned in the animals section followed by an injection of potassium chloride (1-2 mmol/Kg). The tibia bones were harvested and immersed in 10% formaldehyde solution for at least one week. Afterwards, samples were dehydrated in increasing ethanol concentrations (50%, 70%, 100%) for at least 2 days in each solution.

#### Micro-computerized tomography (Micro-CT) analysis

Quantification of bone around the implants was performed using a SkyScan 1272 X-ray Micro-CT scanner (Bruker, USA). Images were acquired at every 0.3° and a resolution of 2016x1334 with a pixel size of 10 μm for a complete 360° rotation. Images were then analyzed using the CT-Analyzer software (CTAn, Bruker). A volume of interest (VOI) was selected around the implants. The NRECON software (Bruker) was used to obtain 3D reconstruction images.

#### Scanning electron microscopy (SEM)

Samples were immersed in ethanol solutions containing increasing concentrations (50%, 70%, 90%, 100%) of methyl-methacrylate resin Technovit 7200 (Kulzer-Heraus, Germany). Samples were then stored in vacuum for 24 h to ensure resin penetration into the tissues, and the resin was photopolymerized using a Histolux light control unit (Kulzer-Heraus) for 24h. The samples were cut in two halves perpendicular to the longitudinal axis of the bone to expose the metallic implants. One of the two halves was polished with 800, 1200 and 4000 SiC abrasive papers and gold-coated by sputtering before visualization in a Phenom XL Desktop SEM. Images were acquired at a working distance of 4 mm and a voltage of 15 kV, and analyzed using the QuPath software^51^. Bone to implant contact (BIC) was calculated using ImageJ as previously described elsewhere^52^.

High resolution images were acquired using a Neon40 Crossbeam FIB-SEM (Carl Zeiss, Germany) at a voltage of 15 kV with a working distance of 8 mm. The percentage of new bone versus total bone was calculated using the QuPath software.

#### Histological staining

Samples were further cut into 500 µm sections with a diamond saw and afterwards polished until 100 µm with SiC abrasive papers. The sections were then stained by Masson’s trichrome staining. Briefly, sections were first stained in Weigert’s hematoxylin for 15 min for staining nuclei and rinsed with tap water for 5 min. Thereafter, sections were stained with Goldner I solution for 7 min and phosphomolybdic acid for 5 min, both for staining connective tissue in red, rinsing with 2% acetic acid after each staining. Finally, sections were stained with Light Green SF solution for 15 min for staining bone in green and rinsed with 2% acetic acid. Sections were then rinsed in tap water and mounted for visualization using an LSM confocal laser scanning microscope (Carl Zeiss, Germany) in stitching mode.

## Acknowledgments

Howard Hughes Medical Institute (D.B.), the Audacious Project (D.L., X.W.), the Defense Threat Reduction Agency grant HDTRA1-21-1-0007 (X.W.), and the DARPA program Harnessing Enzymatic Activity for Lifesaving Remedies (HEALR) under award HR0011-21-2-0012 (B.H.). Spanish Government through project PID2021-125150OB-I00, cofounded by the EU through the European Regional Development Funds (J.M.M., J.G.-M.). Maria Zambrano fellowship funded by European Union-NextGeneration, EU, Ministry of Universities and Recovery, Transformation and Resilience Plan, through a call from Universitat Politècnica de Catalunya (Grant Ref. 2021UPC-MZ-67143, J.G-M). National Institutes of Health (R01 GM109463), Fred Hutchinson Cancer Center (S.K, J.A.C). Predoctoral program AGAUR-FI ajuts (2024 FI-1 00198) Joan Oró, funded by the Secretariat of Universities and Research of the Department of Research and Universities of the Generalitat of Catalonia, and the European Social Plus Fund (D.C.-A.). National Institutes of Health under Grant R35 GM147414 (M.G.C.). Seattle Cancer Consortium Safeway Pilot Awards (funded through the NIH/NCI Cancer Center Support Grant P30 CA015704), the Pew Biomedical Scholars award, and Mike and Debbie Koss (M.G.C). NIH grant R01-HL-131729 (J.L, Y.H, T.S); NIH T90DE021984 grant (Y.T.Z); OFD/BTREC/CTREC Faculty Career Development Fellowship (J.L). Maximizing Investigators’ Research Award (R35GM138036 to C.A.D.) and an Interdisciplinary Training Fellowship (T32CA080416 to I.K.) from the National Institutes of Health; Graduate Research Fellowship (DGE 1762114 to I.K.) from the National Science Foundation. The Generalitat de Catalunya for the ICREA Academia Award (M.-P.G). Electron microscopy data were generated using the Fred Hutchinson Cancer Center Electron Microscopy shared resource, RRID:SCR_022611. The EMSR is supported in part by the Cancer Center Support Grant P30 CA015704-40. We would like to thank Theo Humphreys, Dr. Anvesh Dasari, Steve MacFarlane, and Dr. Caleigh M. Azumaya for their microscopy assistance and knowledge.

Additional acknowledgements

We would also like to thank Longxing Cao, Hua Bai, Brian Coventry for useful discussion and support at the beginning of this project. We would like to thank Natasha Edman, Thomas Schlichthaerle, and the IPD general protein production core for the purified LHD101B-fused C6-79 construct. We thank Lukas Milles and Basile Wicky for providing cloning vectors. We would like to thank Luki Goldschmidt, and Patrick Vecchiato for IT support. We would like to thank the General protein production core at IPD for producing proteins used in animal studies. We would like to thank Inna Goreshnik, and Dionne K. Vafeados in the yeast production core. We would like to thank Madison Kennedy for proofreading the manuscript.

## Author contributions

X.W., J.G-M., S.K, D.L., and D.B. designed the research. X.W and J.G-M designed the initial library. X.W performed the screening experiments and analysis for initial library and affinity maturation. X.W. designed the MPNN-version binder and added a disulfide bond to the designed mini binders. X.W. designed, cloned, expressed, and purified binder constructs for characterization of the binding affinity and specificity to integrin α5β1, cell assay. B.H. designed the ferredoxin scaffold library. J.G-M prepared the covalently linked titanium chips and performed Cell adhesion and spreading assay on the titanium surface. J.G-M. performed real-time PCR and quantified the integrin expressed in cells. S.K. performed the cell attachment and migration inhibition assay and colocalization of binder bound α5β1 with various cellular components. J.G-M and S.K. harvested cells for RNA sequencing experiments. D.L. performed RNA extraction and bulk RNA sequencing and analysis. X.W. and K.A.E.A. performed BLI assays. X.W. and D.L. performed the cell binding assays. Y.L. performed the specificity assay using fluorescence polarization. Y.T.Z. performed the HUVEC tube formation assay. I.K. and C.A.D. designed and performed the hydrogel-based cell encapsulation studies. K.A.E.A. and A.N. prepared integrin α5β1 used for EM and cryo-EM studies. J.L. and K.A.E.A. prepared the integrin α5β1 used for yeast-display assay and BLI. K.A.E.A. and R.W. prepared samples and collected data for EM and cryo-EM. K.A.E.A., R.W., and M.G.C. processed the data and built the cryo-EM model. X.W., R.W, K.A.E.A., A.N. and M.G.C. analyzed the EM and cryo-EM models. A.B.-V. performed rabbit surgical interventions. J.G.-M. and D.C.-A. prepared implant samples and processed the tissues for histomorphometric analyses. X.W, J.G-M wrote the initial manuscript. All authors contributed to the edition and discussion of the manuscript.

## Declaration of generative AI and AI-assisted technologies in the writing process

During the preparation of this work the author(s) used https://chat.openai.com/chat in order to correct grammar. After using this tool/service, the author(s) reviewed and edited the content as needed and take(s) full responsibility for the content of the publication.

## References

1. Koivisto, L., Heino, J., Häkkinen, L. & Larjava, H. Integrins in Wound Healing. Adv. Wound Care 3, 762–783 (2014).

2. Schnittert, J., Bansal, R., Storm, G. & Prakash, J. Integrins in wound healing, fibrosis and tumor stroma: High potential targets for therapeutics and drug delivery. Adv. Drug Deliv. Rev. 129, 37–53 (2018).

3. Kenny, F. N. & Connelly, J. T. Integrin-mediated adhesion and mechano-sensing in cutaneous wound healing. Cell Tissue Res. 360, 571–582 (2015).

4. Tate, M. C. et al. Fibronectin promotes survival and migration of primary neural stem cells transplanted into the traumatically injured mouse brain. Cell Transplant. 11, 283–295 (2002).

5. Harrison, R. P., Medcalf, N. & Rafiq, Q. A. Cell therapy-processing economics: small-scale microfactories as a stepping stone toward large-scale macrofactories. Regen. Med. 13, 159–173 (2018).

6. Patten, J. & Wang, K. Fibronectin in development and wound healing. Adv. Drug Deliv. Rev. 170, 353–368 (2021).

7. Blache, U., Stevens, M. M. & Gentleman, E. Harnessing the secreted extracellular matrix to engineer tissues. Nat Biomed Eng 4, 357–363 (2020).

8. Elmengaard, B., Bechtold, J. E. & Søballe, K. In vivo study of the effect of RGD treatment on bone ongrowth on press-fit titanium alloy implants. Biomaterials 26, 3521–3526 (2005).

9. Elmengaard, B., Bechtold, J. E. & Søballe, K. In vivo effects of RGD-coated titanium implants inserted in two bone-gap models. J. Biomed. Mater. Res. A 75, 249–255 (2005).

10. Li, J. et al. Conformational equilibria and intrinsic affinities define integrin activation. EMBO J. 36, 629–645 (2017).

11. Kapp, T. G. et al. A Comprehensive Evaluation of the Activity and Selectivity Profile of Ligands for RGD-binding Integrins. Sci. Rep. 7, 1–13 (2017).

12. Xia, W. & Springer, T. A. Metal ion and ligand binding of integrin α5β1. Proc. Natl. Acad. Sci. U. S. A. 111, 17863–17868 (2014).

13. Ludwig, B. S., Kessler, H., Kossatz, S. & Reuning, U. RGD-Binding Integrins Revisited: How Recently Discovered Functions and Novel Synthetic Ligands (Re-)Shape an Ever-Evolving Field. Cancers 13, (2021).

14. Marsico, G., Russo, L., Quondamatteo, F. & Pandit, A. Glycosylation and Integrin Regulation in Cancer. Trends Cancer Res. 4, 537–552 (2018).

15. Takagi, J., Strokovich, K., Springer, T. A. & Walz, T. Structure of integrin alpha5beta1 in complex with fibronectin. EMBO J. 22, 4607–4615 (2003).

16. Schumacher, S. et al. Structural insights into integrin α5β1 opening by fibronectin ligand. Sci Adv 7, (2021).

17. Roy, A. et al. De novo design of highly selective miniprotein inhibitors of integrins αvβ6 and αvβ8. Nat. Commun. 14, 5660 (2023).

18. Cao, L. et al. Design of protein-binding proteins from the target structure alone. Nature 605, 551–560 (2022).

19. Cao, L. et al. De novo design of picomolar SARS-CoV-2 miniprotein inhibitors. Science 370, 426–431 (2020).

20. Hiranuma, N. et al. Improved protein structure refinement guided by deep learning based accuracy estimation. Nat. Commun. 12, 1340 (2021).

21. Kumar, S., Stainer, A., Dubrulle, J., Simpkins, C. & Cooper, J. A. Cas phosphorylation regulates focal adhesion assembly. Elife 12, (2023).

22. Lambert, A. W., Ozturk, S. & Thiagalingam, S. Integrin signaling in mammary epithelial cells and breast cancer. ISRN Oncol. 2012, 493283 (2012).

23. Ly, T. et al. Proteome-wide analysis of protein abundance and turnover remodelling during oncogenic transformation of human breast epithelial cells. Wellcome Open Res 3, 51 (2018).

24. Moore, K. M. et al. Therapeutic targeting of integrin αvβ6 in breast cancer. J. Natl. Cancer Inst. 106, (2014).

25. Gilcrease, M. Z. Integrin signaling in epithelial cells. Cancer Lett. 247, 1–25 (2007).

26. Anderson, J. M., Li, J. & Springer, T. A. Regulation of integrin α5β1 conformational states and intrinsic affinities by metal ions and the ADMIDAS. Mol. Biol. Cell 33, ar56 (2022).

27. Nagae, M. et al. Crystal structure of α5β1 integrin ectodomain: atomic details of the fibronectin receptor. J. Cell Biol. 197, 131–140 (2012).

28. Humphries, J. D. et al. Molecular basis of ligand recognition by integrin alpha5beta 1. II. Specificity of arg-gly-Asp binding is determined by Trp157 OF THE alpha subunit. J. Biol. Chem. 275, 20337–20345 (2000).

29. De Franceschi, N., Hamidi, H., Alanko, J., Sahgal, P. & Ivaska, J. Integrin traffic - the update. J. Cell Sci. 128, 839–852 (2015).

30. Sottile, J. & Chandler, J. Fibronectin matrix turnover occurs through a caveolin-1-dependent process. Mol. Biol. Cell 16, 757–768 (2005).

31. McCall, J. D. & Anseth, K. S. Thiol–Ene Photopolymerizations Provide a Facile Method To Encapsulate Proteins and Maintain Their Bioactivity. Biomacromolecules 13, 2410–2417 (2012).

32. McCall, J. D., Luoma, J. E. & Anseth, K. S. Covalently tethered transforming growth factor beta in PEG hydrogels promotes chondrogenic differentiation of encapsulated human mesenchymal stem cells. Drug Deliv. Transl. Res. 2, 305–312 (2012).

33. Sridhar, B. V., Doyle, N. R., Randolph, M. A. & Anseth, K. S. Covalently tethered TGF-β1 with encapsulated chondrocytes in a PEG hydrogel system enhances extracellular matrix production. J. Biomed. Mater. Res. A 102, 4464–4472 (2014).

34. Walters, B. et al. Engineering the geometrical shape of mesenchymal stromal cells through defined cyclic stretch regimens. Sci. Rep. 7, 6640 (2017).

35. Lichtman, M. K., Otero-Vinas, M. & Falanga, V. Transforming growth factor beta (TGF-β) isoforms in wound healing and fibrosis. Wound Repair Regen. 24, 215–222 (2016).

36. Wu, M., Chen, G. & Li, Y.-P. TGF-β and BMP signaling in osteoblast, skeletal development, and bone formation, homeostasis and disease. Bone Res 4, 16009 (2016).

37. Takagi, J. & Springer, T. A. Integrin activation and structural rearrangement. Immunol. Rev. 186, 141–163 (2002).

38. Wicky, B. I. M. et al. Hallucinating symmetric protein assemblies. Science 378, 56–61 (2022).

39. Ohi, M., Li, Y., Cheng, Y. & Walz, T. Negative Staining and Image Classification - Powerful Tools in Modern Electron Microscopy. Biol. Proced. Online 6, 23–34 (2004).

40. Suloway, C. et al. Automated molecular microscopy: the new Leginon system. J. Struct. Biol. 151, 41–60 (2005).

41. Zivanov, J. et al. New tools for automated high-resolution cryo-EM structure determination in RELION-3. Elife 7, (2018).

42. Punjani, A., Rubinstein, J. L., Fleet, D. J. & Brubaker, M. A. cryoSPARC: algorithms for rapid unsupervised cryo-EM structure determination. Nat. Methods 14, 290–296 (2017).

43. Mastronarde, D. N. Automated electron microscope tomography using robust prediction of specimen movements. J. Struct. Biol. 152, 36–51 (2005).

44. Zheng, S. Q. et al. MotionCor2: anisotropic correction of beam-induced motion for improved cryo-electron microscopy. Nat. Methods 14, 331–332 (2017).

45. Emsley, P. & Cowtan, K. Coot: model-building tools for molecular graphics. Acta Crystallogr. D Biol. Crystallogr. 60, 2126–2132 (2004).

46. Croll, T. I. ISOLDE: a physically realistic environment for model building into low-resolution electron-density maps. Acta Crystallogr D Struct Biol 74, 519–530 (2018).

47. Guillem-Marti, J. et al. RGD Mutation of the Heparin Binding II Fragment of Fibronectin for Guiding Mesenchymal Stem Cell Behavior on Titanium Surfaces. ACS Appl. Mater. Interfaces 11, 3666–3678 (2019).

48. Heras-Parets, A., Ginebra, M.-P., Manero, J. M. & Guillem-Marti, J. Guiding fibroblast activation using an RGD-mutated heparin binding II fragment of fibronectin for gingival titanium integration. Adv. Healthc. Mater. 12, e2203307 (2023).

49. Zhao, Y. T. et al. F-domain valency determines outcome of signaling through the angiopoietin pathway. EMBO Rep. 22, e53471 (2021).

50. Stirling, D. R. et al. CellProfiler 4: improvements in speed, utility and usability. BMC Bioinformatics 22, 433 (2021).

51. Bankhead, P. et al. QuPath: Open source software for digital pathology image analysis. Sci. Rep. 7, 16878 (2017).

52. Manresa, C., Bosch, M., Manzanares, M. C., Carvalho, P. & Echeverría, J. J. A new standardized-automatic method for bone-to-implant contact histomorphometric analysis based on backscattered scanning electron microscopy images. Clin. Oral Implants Res. 25, 702–706 (2014).

