## Supplemental for "De Novo Design of Integrin α5β1 Modulating Proteins for Regenerative Medicine"

**Supplemental items**

Figure S1 Experimental screening of α5β1 binders, related to Figure 1.

Figure S2 Competition curves, and calculated $K_D$ values from fluorescence anisotropy, related to Figure 2.

Figure S3 Colocalization of NeoNectin with ITGB1 and endosomal markers in MCF10A cells, related to Figure 2.

Figure S4 Cryo-EM data processing schematic, related to Figure 3.

Figure S5 Negative stain of integrin α5β1 alone and in complex with NeoNectin and NeoNectin variants, related to Figure 3.

Figure S6 Soluble NeoNectin inhibits α5β1-mediated cellular behaviors, related to Figure 4.

Figure S7 Hydrogel modification enhances cells spreading, related to Figure 5.

Figure S8 Modulation of cell adhesion by NeoNectin immobilized onto Ti discs, related to Figure 5.

Figure S9 Scatterplots of gene expression against bare Ti discs for FN-, NeoNectin-, and RGD-grafted Titanium discs, related to Figure 5.

Figure S10: NeoNectin-grafted titanium implant outperforms FN- and RGD- grafted, and bare titanium implants in stimulating implant integration and bone growth, related to Figure 6.

Table S1. Cryo-EM Data Collection and Processing Statistics

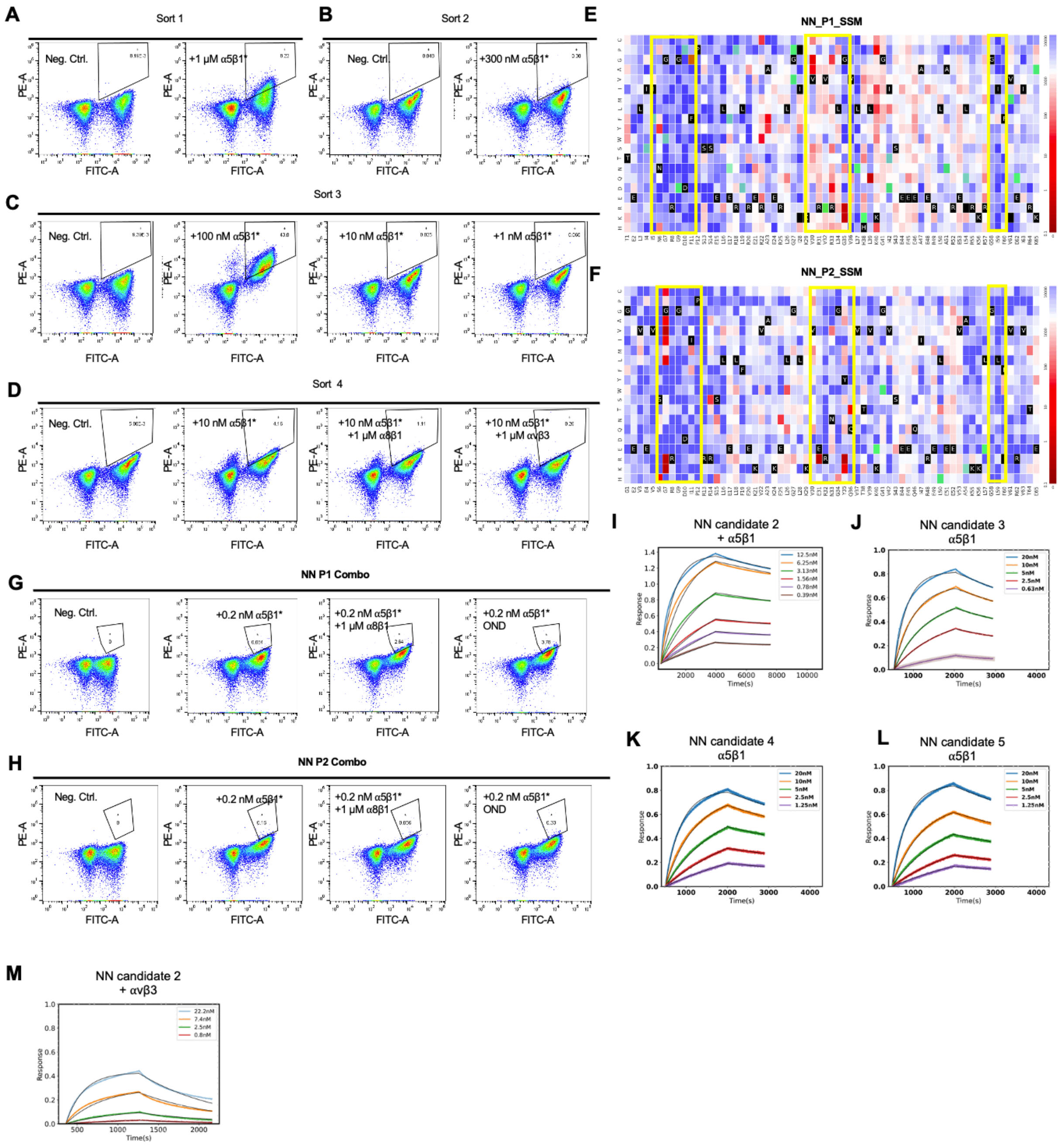


**Figure S1 Experimental screening of α5β1 binders, related to Figure 1**.

(A-D) Yeast cells displaying the SSM library of integrin α5β1 binders were incubated with various concentrations of biotinylated α5β1 in the presence or absence of non-labelled αvβ3 or α8β1. Labeled α5β1 binding to cells (y axis) was monitored with flow cytometry.

(E-F) SSM analysis of NN parent design 1 (NN P1)and parent design 2 (NN P2). Loop1, loop3 and loop5 were highlighted. The affinity of each variant was colored by red to white to blue gradient: red indicates the tightest binders, blue the weakest binders, and green indicates variants without sufficient data for analysis.

(G-H) Combinatorial libraries of the mutations that improved binding affinity, as identified by the red positions in Figure E and F.

(I-L) BLI binding affinity traces for NeoNectin candidates against α5β1 in the resting buffer (20 mM Tris, pH = 7.4, 1 mM $Ca^{2+}$, 1 mM $Mg^{2+}$). From left to right, $K_D$= 0.4, 1.5, 1.3, and 1.3 nM respectively.

(M) BLI binding affinity traces for NeoNectin candidate 2 against integrin αvβ3 in the resting buffer. $K_D$= 6.1 nM

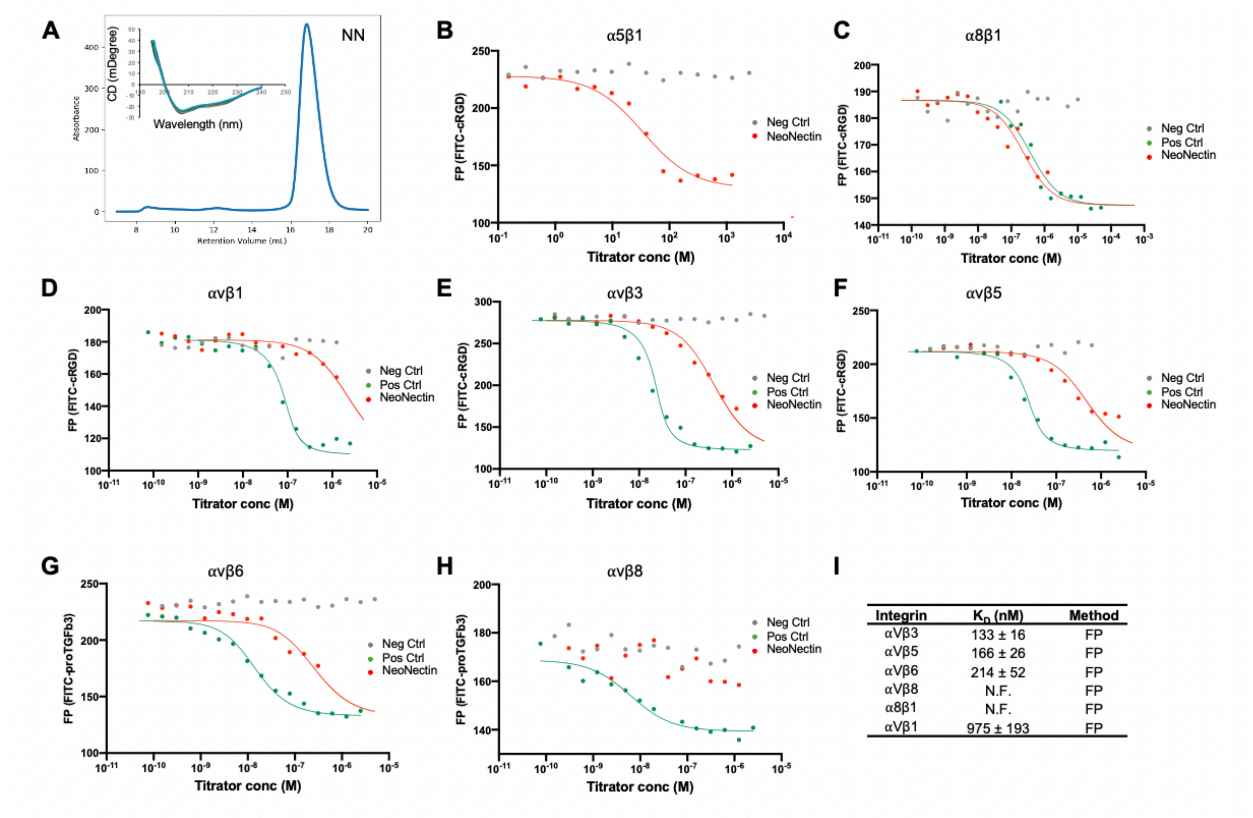


**Figure S2 Competition curves, and calculated $K_D$ values from fluorescence anisotropy, related to Figure 2.**

(A) Size Exclusion Chromatography trace of NeoNectin purified from 40 milliliter (ml) media and Circular dichroism spectra of NeoNectin at different temperatures (navy, 25 °C; orange, 55 °C; green, 75 °C, blue 95 °C).

(B) Binding affinity of NeoNectin to integrin α5β1 was measured by competing 10 nM FITC-cyclic-RGD binding to 100 nM α5β1. Neg Ctrl: αvβ3_ab7. $K_D$ couldn't be fitted.

(C) Binding affinity of NeoNectin to integrin α8β1 was measured by competing 10 nM FITC-cyclic-RGD binding to 1000 nM α8β1. Neg Ctrl: αvβ3_ab13; Pos Ctrl: cRGD

(D) Binding affinity of NeoNectin to integrin αvβ1 was measured by competing 10 nM FITC-cyclic-RGD binding to 200 nM αvβ1. Neg Ctrl: αvβ3_ab13; Pos Ctrl: αvβ1_ab5

(E) Binding affinity of NeoNectin to integrin αvβ3 was measured by competing 10 nM FITC-cyclic-RGD binding to 50 nM αvβ3. Neg Ctrl: αvβ6_ab4; Pos Ctrl: αvβ3_ab13

(F) Binding affinity of NeoNectin to integrin αvβ5 was measured by competing 10 nM FITC-cyclic-RGD binding to 50 nM αvβ5. Neg Ctrl: αvβ3_ab13; Pos Ctrl: αvβ5_ab9

(G) Binding affinity of NeoNectin to integrin αvβ6 was measured by competing 10 nM FFITC-proTGFβ3 binding to 10 nM αvβ6. Neg Ctrl:αvβ3_ab7; Pos Ctrl: αvβ6_ab6

(H) Binding affinity of NeoNectin to integrin αvβ8 was measured by competing 10 nM FFITC-proTGFβ3 binding to 250 nM αvβ8. Neg Ctrl:αvβ3_ab13 ; Pos Ctrl: αvβ8_ab4 (dual specific to αvβ6 and αvβ8).

(I) Binding affinity of NeoNectin to the soluble ectodomains of RGD-binding integrins by fluorescent polarization competitive binding assays from C to H.

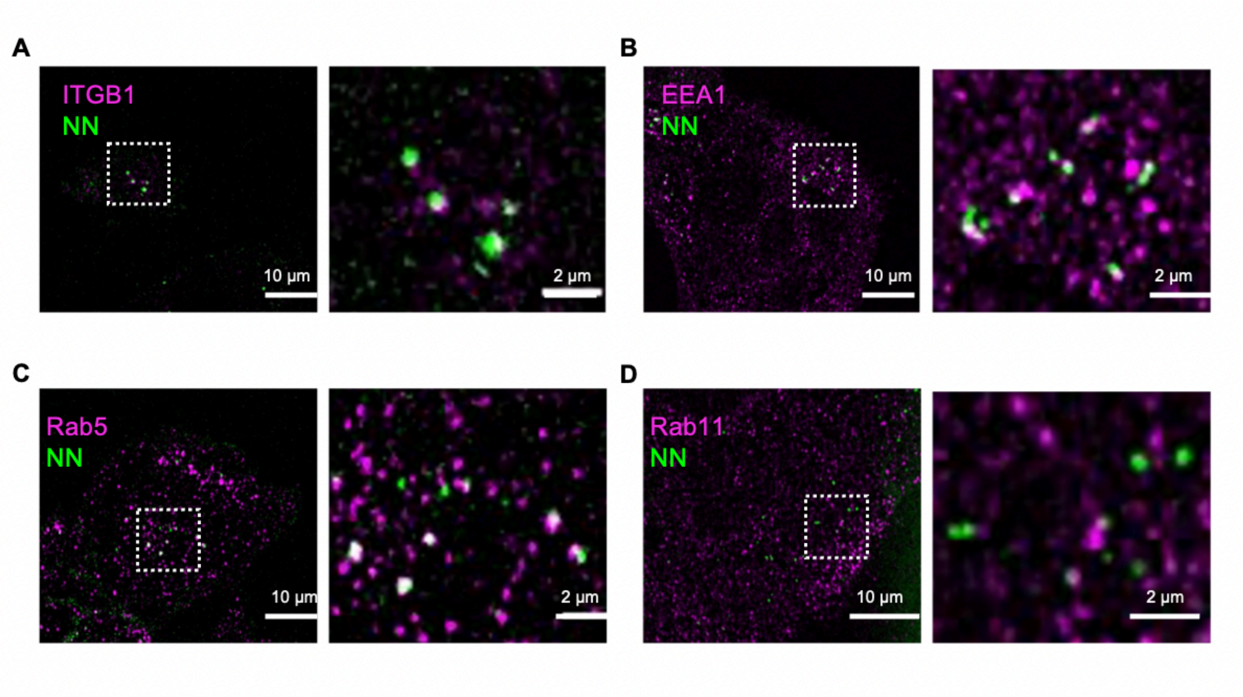


**Figure S3 Colocalization of NeoNectin with ITGB1 and endosomal markers in MCF10A cells, related to Figure 2.**

(A-D) Cross-section confocal images showing colocalization of NeoNectin and integrin β1 subunit (A), NeoNectin and EEA1 (B), NeoNectin and Rab5 (C), and NeoNectin and Rab11. Right panel is a zoomed view of the left panel. Scale bars are 10 µm and 2 µm, respectively.

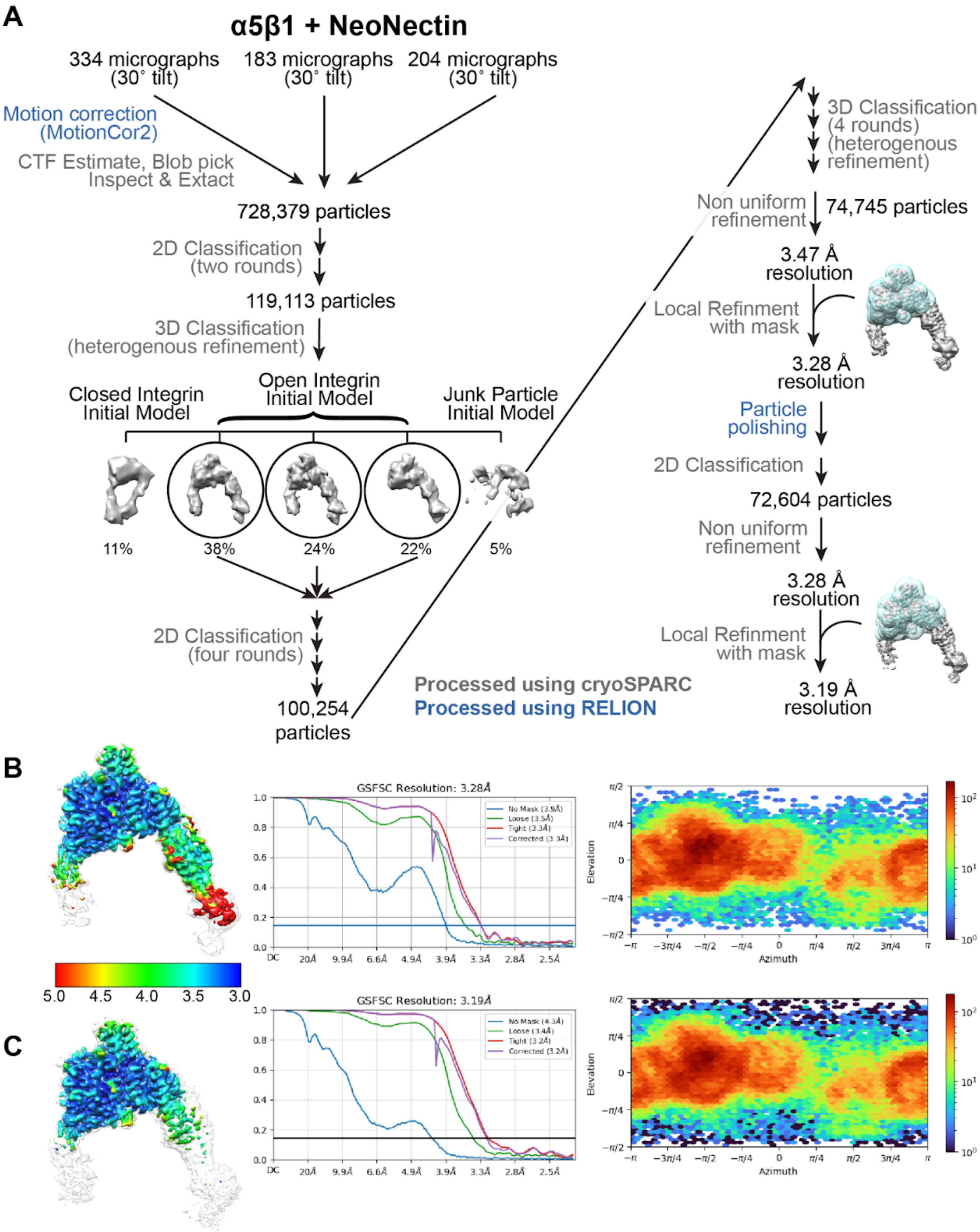


**Figure S4 Cryo-EM data processing schematic, related to Figure 3.**

(A) An overview summarizing cryo-EM data processing for the α5β1+NeoNectin complex. Particle numbers at key steps and percentages for each class are indicated.

(B) Left: Sharpened map of α5β1+NeoNectin complex colored based on a local resolution estimate, unsharpened map shown in semi-transparent white at a lower threshold. Middle: Gold-standard Fourier shell correlation plot. Right: Orientational distribution plot of the α5β1 in complex with NeoNectin.

(C) Left: Final local refined and sharpened map of α5β1+NeoNectin complex colored based on a local resolution estimate, unsharpened map shown in semi-transparent white at a lower threshold. Middle: Gold-standard Fourier shell correlation plot. Right: Orientational distribution plot of the locally-refined α5β1 in complex with NeoNectin.

A

B

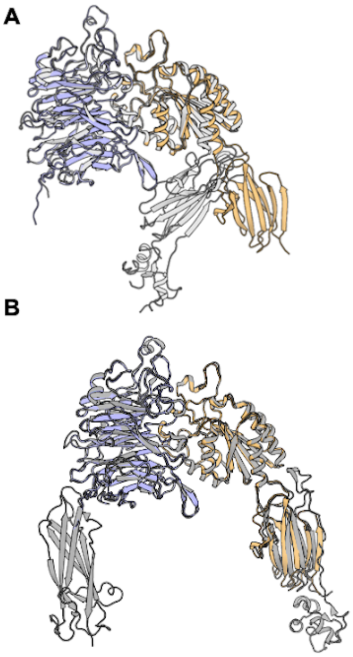

C

| Loop | NeoNectin | a5 | B1 | Type of Interaction |
|---|---|---|---|---|
| L1 | Arg 8 | Phe 187 | | Shape Complementarity |
| L1 | Arg 8 | Gln 189 | | Shape Complementarity |
| L1 | Arg 8 | Gln 221 | | Shape Complementarity |
| L1 | Arg 8 | Ser 224 | | Hydrogen Bond/ Shape Complementarity |
| L1 | Arg 8 | Asp227 | | Salt Bridge / Hydrogen Bond |
| L1 | Gly 9 | Phe 187 | | Shape Complementarity |
| L3 | Asp 33 | Glycan 275 | | Shape Complementarity |
| L3 | His 34 | Ser 224 | | Shape Complementarity |
| L3 | His 34 | Ile 225 | | Shape Complementarity |
| L5 | Trp 60 | Ser 156 | | Shape Complementarity |
| L5 | Trp 60 | Trp 157 | | Shape Complementarity |
| L5 | Trp 60 | Ala 159 | | Shape Complementarity |
| L1 | Gly 9 | | Leu 225 | Shape Complementarity |
| L1 | Asp 10 | | Ser 132 | MIDAS Ion Coordination |
| L1 | Asp 10 | | Tyr 133 | Shape Complementarity |
| L1 | Asp 10 | | Ser 134 | MIDAS Ion Coordination |
| L1 | Asp 10 | | Asn 224 | Shape Complementarity |
| L1 | Asp 10 | | Asp 226 | Shape Complementarity |
| L1 | Asp 10 | | Ser 227 | Shape Complementarity |
| L1 | Asp 10 | | Glu 229 | MIDAS Ion Coordination |
| L1 | Phe 11 | | Tyr 133 | Shape Complementarity |
| L1 | Pro 12 | | Tyr 133 | Shape Complementarity |
| none | Ser 14 | | Asp 137 | Shape Complementarity |
| L3 | Glu 15 | | Lys 136 | Hydrogen Bond |
| L3 | His 34 | | Glu 320 | Salt Bridge |
| L3 | Lys 35 | | Glu 320 | Salt Bridge |
| L5 | Gly 58 | | Thr 188 | Shape Complementarity |

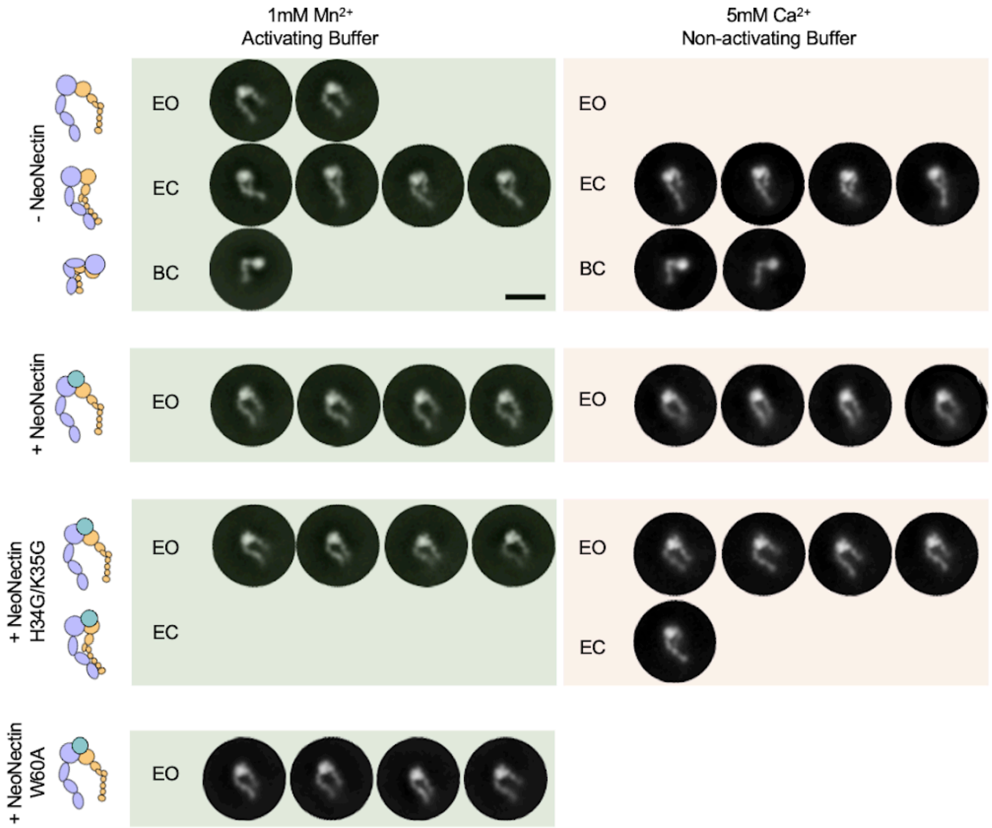

**Figure S5 Negative stain of integrin α5β1 alone and in complex with NeoNectin and NeoNectin variants, related to Figure 3.**

(A) Overlay of α5β1 in complex with NeoNectin (lavender, light orange) and α5β1 in complex with FN (gray, PDB:7NWL). The secondary structural elements of β-propeller and βI domain overlay with very high similarity, and we attribute the slight shift in the hybrid domain to flexibility, as both models were built into maps with anisotropic resolution.

(B) Overlay of α5β1 in complex with NeoNectin (lavender, light orange) and α5β1 in complex with RGD peptide (white, PDB:4WK2).

(C) Summary of interactions between NeoNectin and α5β1.

(D-F) Representative 2D negative stain class averages of α5β1 in activating (1 mM $Mn^{2+}$) and non-activating (5 mM $Ca^{2+}$) buffer conditions. Integrins are categorized into the three canonical conformations: extended open (EO), extended closed (EC) and bent closed (BC). The number of classes shown is representative of the number of total particles in that conformation. (D) α5β1 alone. (E) α5β1 in complex with NeoNectin. (F) α5β1 in complex with the NeoNectin H34G/K35G variant. We note that there is not a clear density that can be attributed to NeoNectin variant in the EC conformation, and thus this class could represent apo integrins.

(G) Representative 2D negative stain class averages of α5β1 bound to NeoNectin W60A variant in activating buffer (1 mM $Mn^{2+}$). **Figure S6 Soluble NeoNectin inhibits α5β1-mediated cellular behaviors, related to Figure 4.**

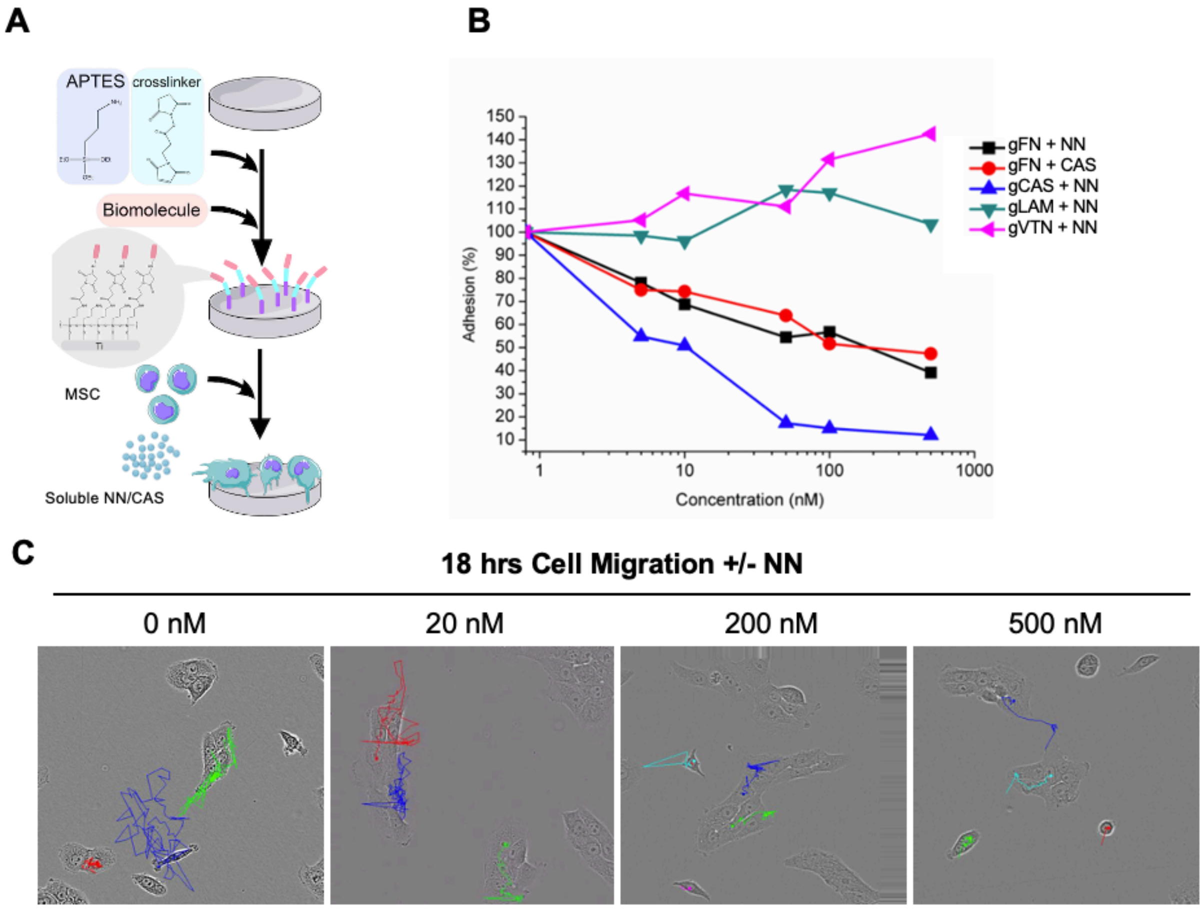


(A) Schematic of the covalent immobilization of NeoNectin on Ti discs.

(B) Percentage of cells adhered on titanium discs Grafted with FN (gFN), the cell attachment fragment from FN (gCAS), laminin (gLAM), or vitronectin (gVTN) when co-incubated with soluble NeoNectin (NN) or soluble CAS .

(C) Trajectories of individual cells tracked over an 18 hours imaging period in presence of 0, 20, 200, and 500nM NeoNectin.

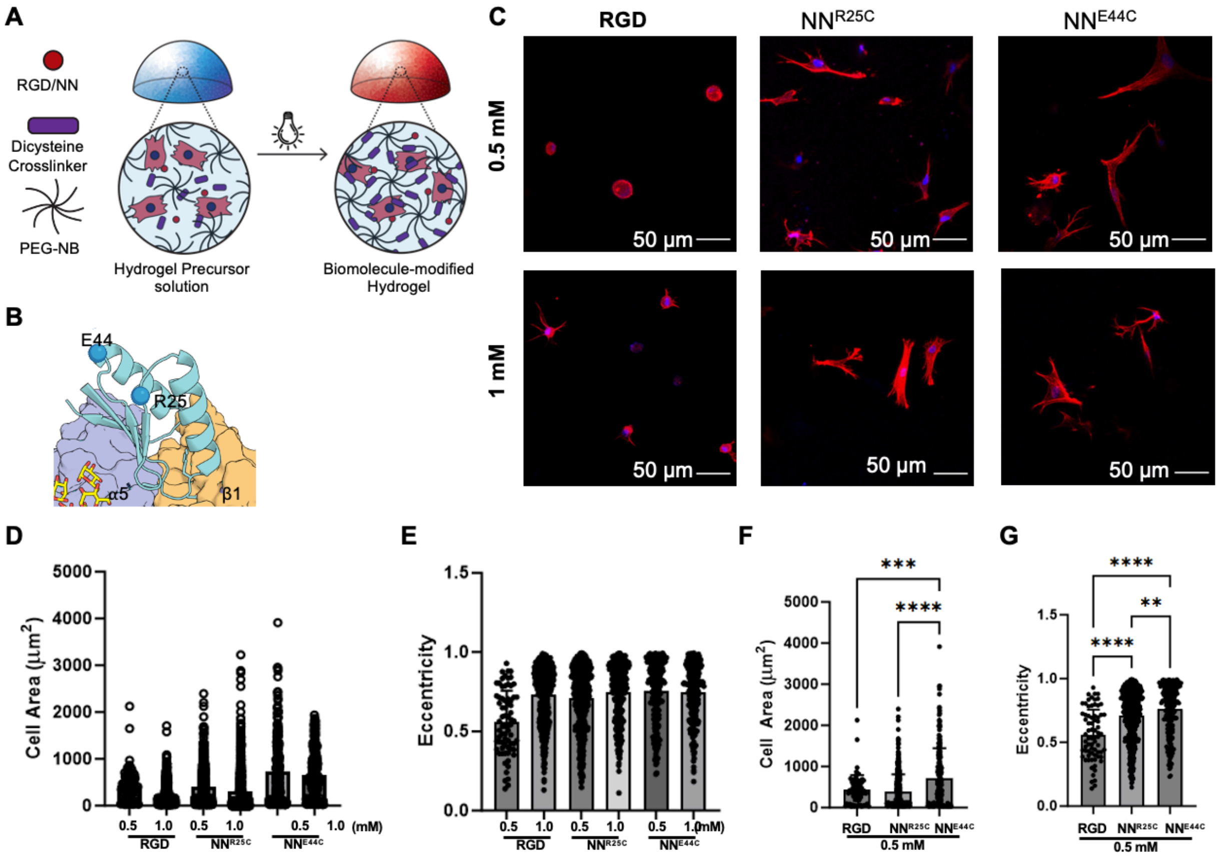


**Figure S7 Hydrogel modification enhances cells spreading, related to Figure 5.**

(A) Schematic representation of the radical thiol-ene-based covalent immobilization of NeoNectin in hydrogel.

(B) NeoNectin variants with cysteine mutations.

(C) Representative images of MSCs encapsulated in RGD, $NN^{R25C}$, and $NN^{E44C}$ modified hydrogels post 5d of culture.

(D-E) Quantification of cell spread area and eccentricity across all conditions tested.

F-G, Quantification of cell area and eccentricity in the 0.5 mM conditions, which displayed the greatest variation across all tested. Error bars = S.D. One Way ANOVA, Tukey's Post-hoc Test. ** = $p < 0.01$, *** = $p < 0.001$, **** = $p < 0.0001$.

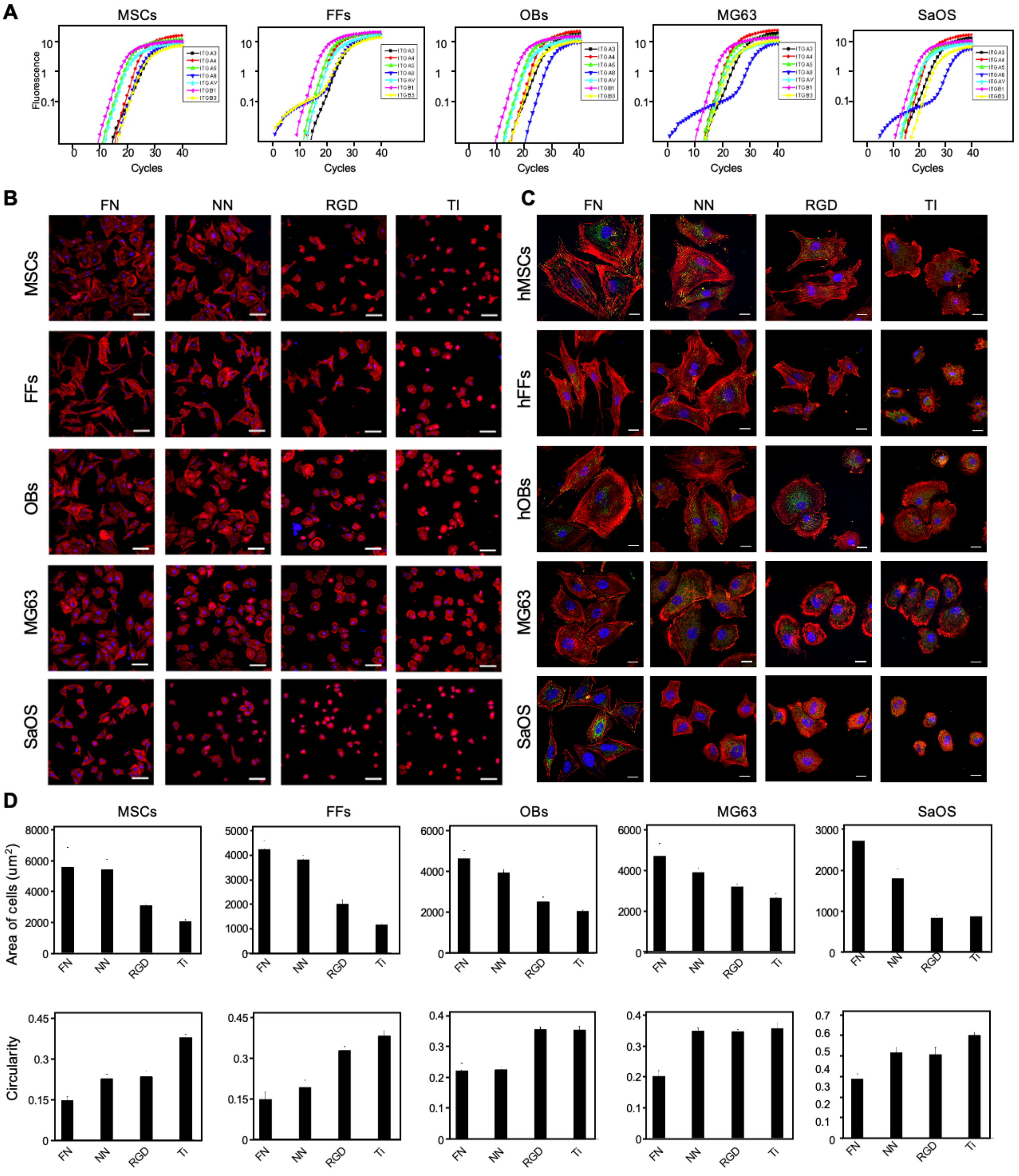


**Figure S8 Modulation of cell adhesion by NeoNectin immobilized onto Ti discs, related to Figure 5.**

(A) Real-time PCR amplification plots showing the expression of different integrins in MSCs, FFs, OBs, MG63 and SaOS-2 cells.

(B) Representative immunofluorescence images of MSCs, FFs, OBs, MG63 and SaOS-2 cells after 4h of adhesion on the different functionalized Ti discs. The scale bar denotes 100 μm.

(C) Representative high magnification images from B. The scale bar denotes 10 μm.

(D) Quantification of the area and circularity of cells from B.

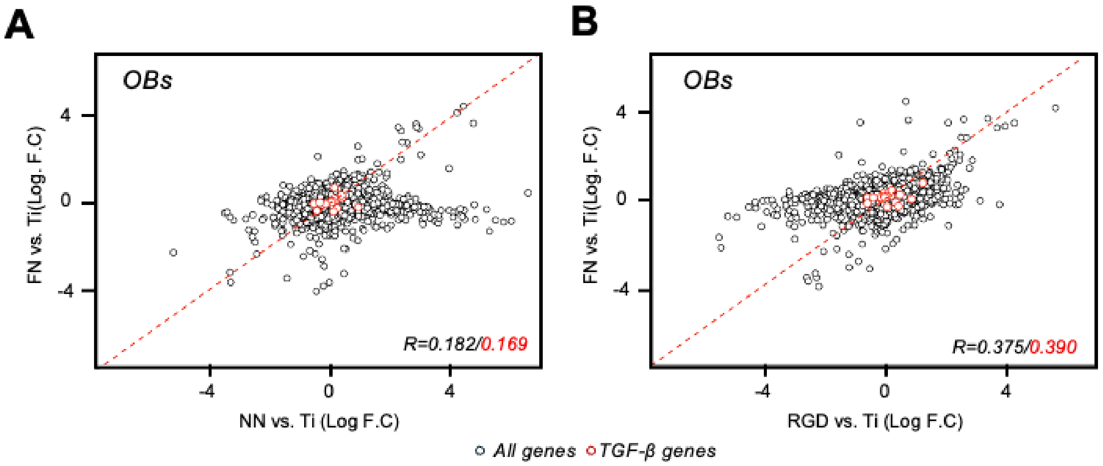


**Figure S9 Scatterplots of gene expression against bare Ti discs for FN-, NeoNectin-, and RGD-grafted Titanium discs, related to Figure 5.**

(A) Scatterplots of gene expression against bare Ti discs for FN-, and NeoNectin-grafted titanium discs. The genes of the TGF-β pathway for each cell type are highlighted in red circles.

(B) Scatterplots of gene expression against bare Ti discs for FN-, and RGD-grafted titanium discs. The genes of the TGF-β pathway for each cell type are highlighted in red circles.

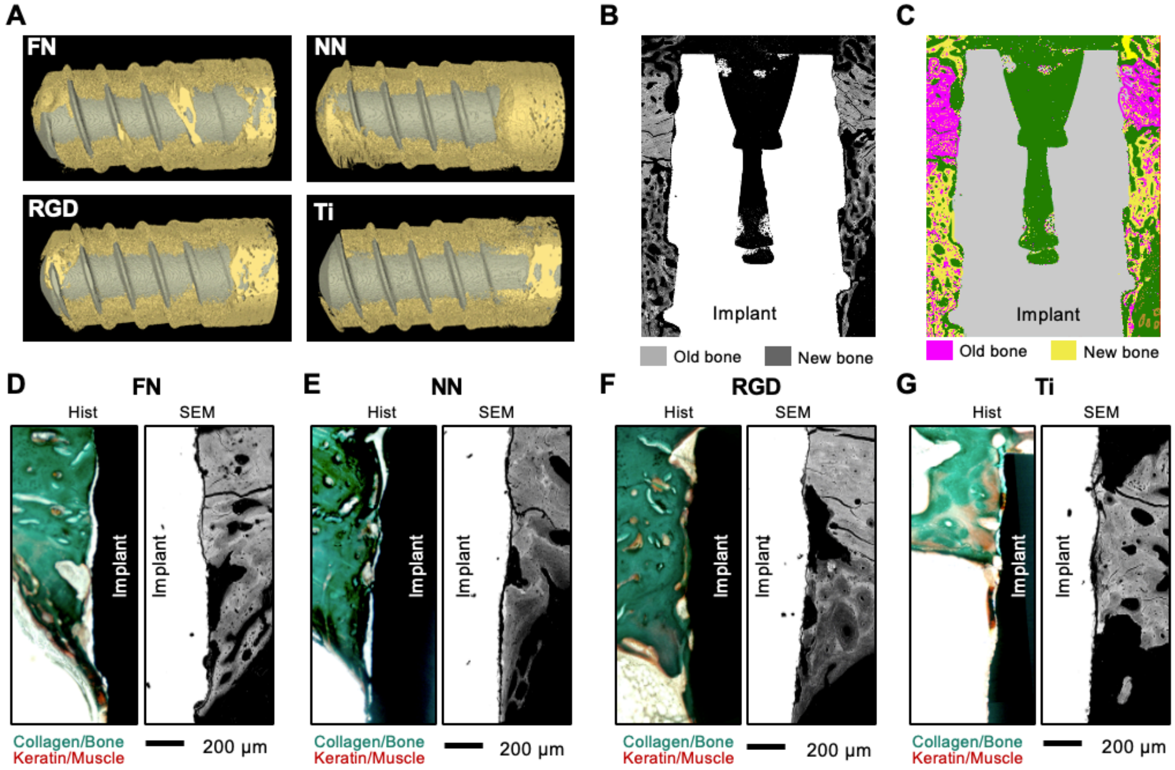


**Figure S10: NeoNectin-grafted titanium implant outperforms FN- and RGD- grafted, and bare titanium implants in stimulating implant integration and bone growth, related to Figure 6.**

(A) Representative micro-CT 3D reconstruction images showing bone (yellow) around the grafted or bare titanium implants (gray) 6 weeks post-surgery.

(B-C) Example of image processing for the calculation of new bone percentage. SEM images showing old bone in light gray and new bone in dark gray (B) were processed to show each bone type as pink and yellow, respectively.

(D-G) Representative histological staining (left) and SEM (right) images of longitudinal sections 6 weeks post-implantation showing the implants conjugated with indicated molecules inserted into the tibia of rabbits.

**Table S1. Cryo-EM Data Collection and Processing Statistics**

| | NeoNectin:α5β1 (EMDB-(TBD)) (PDB (TBD)) |
|---|---|
| **Data collection and processing** | |
| **Magnification** | 36,000x |
| **Voltage (kV)** | 200 |
| **Electron exposure (e–/Å$^2$)** | 50 |
| **Defocus range (μm)** | |
| **Pixel size (Å)** | 1.122 |
| **Symmetry imposed** | C1 |
| **Initial particle images (no.)** | 372,064 |
| **Final particle images (no.)** | 72,604 |
| **Map resolution (Å)** | 3.28 (overall); 3.19 (local refinement) |
| **FSC threshold** | 0.143 |
| **Map resolution range (Å)** | 2.92- 17.44 (overall); 2.92-9.65 (local refinement) |
